# Crosstalk between the tRNA methyltransferase Trm1 and RNA chaperone La influences eukaryotic tRNA maturation

**DOI:** 10.1101/2023.04.12.536578

**Authors:** Jennifer Porat, Ana Vakiloroayaei, Brittney M. Remnant, Mohammadaref Talebi, Taylor Cargill, Mark A. Bayfield

## Abstract

tRNAs undergo an extensive maturation process involving post-transcriptional modifications often associated with tRNA structural stability and promoting the native fold. Impaired post-transcriptional modification has been linked to human disease, likely through defects in translation, mitochondrial function, and increased susceptibility to degradation by various tRNA decay pathways. More recently, evidence has emerged that bacterial tRNA modification enzymes can act as tRNA chaperones to guide tRNA folding in a manner independent from catalytic activity. Here, we provide evidence that the fission yeast tRNA methyltransferase Trm1, which dimethylates nuclear– and mitochondrial-encoded tRNAs at G26, can also promote tRNA functionality in the absence of catalysis. We show that wild type and catalytic-dead Trm1 are active in an *in vivo* tRNA-mediated suppression assay and possess RNA strand annealing and dissociation activity *in vitro*, similar to previously characterized RNA chaperones. Trm1 and the RNA chaperone La have previously been proposed to function synergistically in promoting tRNA maturation, yet we surprisingly demonstrate that La binding to nascent pre-tRNAs decreases Trm1 tRNA dimethylation *in vivo* and *in vitro*. Collectively, these results support the hypothesis for tRNA modification enzymes that combine catalytic and non-catalytic activities to promote tRNA maturation, as well as expand our understanding of how La function can influence tRNA modification.

## Introduction

Owing to their critical role in translation, tRNAs are subject to numerous processing and quality control steps to ensure their structural stability and functionality. In eukaryotes, this involves removal of a 5’ leader and 3’ trailer sequence; where applicable, removal of an intron; CCA addition and aminoacylation of the mature 3’ end; as well as the acquisition of post-transcriptional modifications (reviewed in (1, 2)). While most post-transcriptional modifications are non-essential, especially in yeast, the combination of modifications on a given tRNA is an important determinant for structural stability and functionality. Modifications to the anticodon loop affect translational fidelity by maintaining the correct open reading frame and influencing codon-anticodon base-pairing, whereas modifications to the tRNA body primarily affect structure and folding (3). Importantly, studies have found that pre-tRNAs lacking certain post-transcriptional modifications are prone to misfolding and subsequently targeted for decay by the nuclear surveillance machinery (4, 5), while hypomodified mature tRNAs are degraded by the rapid tRNA decay pathway (6–8).

Links between pre-tRNA structure and escape from decay are especially relevant in the context of the function of the eukaryotic RNA chaperone La. The La protein interacts with the 3’ uridylate trailer of RNA Polymerase III transcripts, including nascent pre-tRNAs, through a binding pocket formed by the eponymous La motif and RNA recognition motif 1 (RRM1) (9–12). As such, a major role of the La protein is to protect the 3’ end of pre-tRNAs from exonucleolytic degradation and direct the order of pre-tRNA end processing (reviewed in (13)). La binding to the 3’ trailer results in 5’ leader processing by RNase P followed by 3’ trailer cleavage, the latter of which enables La dissociation and recycling onto a new pre-tRNA substrate (14, 15). 3’ end binding by La is particularly important for structurally defective pre-tRNAs, which rely on La for protection from decay by the nuclear exosome (16). However, 3’ end protection alone is insufficient to rescue increasingly defective pre-tRNAs from nuclear surveillance, necessitating a second activity mapping to the canonical RNA binding surface of the RRM1 (15, 16). Further insight into this alternate activity revealed that the RRM1 interacts with the pre-tRNA body, where it can act as an RNA chaperone to assist pre-tRNA folding (15, 17, 18). Coupling of La’s two distinct binding modes—3’ uridylate binding and contacts to the tRNA body—enables high-affinity engagement of pre-tRNAs, resulting in their stabilization and proper folding (15).

As tRNAs can also achieve their native, functional conformation through post-transcriptional modifications, the La protein has been proposed to function redundantly with tRNA modification enzymes. In agreement with this, deletion of La is synthetically lethal with the deletion of several tRNA modification enzymes in budding and fission yeast grown at elevated temperatures, where tRNA misfolding can occur (18, 19). Rescue of synthetic lethality by wild type La, but not RRM1 mutants linked to defective RNA chaperone function, further supports the idea that La and tRNA modification enzymes have roles in promoting correct tRNA folding (18). Among the modification enzymes that function redundantly with La, N2, N2-dimethylation at G26 by the tRNA methyltransferase Trm1 (20), has been demonstrated to stabilize pre-tRNA *in vitro* and *in vivo* (18, 21). G26 dimethylation has been linked to structural stability at the junction between the anticodon and variable arm, or hinge region, in that lack of G26 dimethylation results in increased accessibility of nucleotides in the hinge of a pre-tRNA substrate to chemical probing by lead acetate *in vitro* (18). While studies on the structure stabilizing effect of Trm1-catalyzed methylation have largely been limited to pre-tRNAs in the nucleus, nuclear-encoded Trm1 contains an N-terminal mitochondrial targeting sequence, leading to the modification of select G26-containing mitochondrial-encoded tRNAs (22, 23). The human mitochondrial-localized Trm1 has been linked to promoting protein synthesis, cellular proliferation, and redox homeostasis through modification of mt-tRNA Ile^UAU^, mt-tRNA Ala^UGC^, and mt-tRNA Arg^UCG^ (22). In contrast, budding yeast produce 2 Trm1 isoforms arising from alternate transcription start sites that yield proteins differing in the presence of the mitochondrial targeting sequence (23). While the additional N-terminal sequence enhances the efficiency of mitochondrial targeting, it is not strictly required for mitochondrial import, as both isoforms are capable of methylating mitochondrial-encoded tRNAs (23).

The structural and functional importance of post-transcriptional modifications on both nuclear– and mitochondrial-encoded tRNAs has been well-established (reviewed in (24)), but recent evidence points to the idea that post-transcriptional modification enzymes may have alternate functions related to tRNA folding (2). Notably, the bacterial tRNA pseudouridine synthase TruB and methyltransferase TrmA promote tRNA folding even in the absence of catalysis, leading to their characterization as tRNA chaperones (25, 26). Although a catalytically inactive mutant of the eukaryotic TrmA homolog Trm2 can rescue a growth defect associated with a mutant allele of tRNA Ser^CGA^, suggesting that the dual function of tRNA modification enzymes is evolutionarily conserved (27), to date there has been no mechanistic insight into how eukaryotic tRNA modification enzymes promote tRNA folding independent from catalytic activity.

Here, we investigated the importance of catalytic versus non-catalytic functions of the tRNA methyltransferase Trm1 from *Schizosaccharomyces pombe* and examined crosstalk between Trm1 and function of the *S. pombe* La homolog Sla1, an established RNA chaperone. We confirmed that similar to budding yeast, an N-terminal mitochondrial localization signal enabled select modification of mitochondrial-encoded, but not nuclear-encoded tRNAs. We demonstrated that mutation of a key catalytic residue resulted in a complete loss of Trm1 modification at G26 *in vitro* and *in vivo*, but that this mutant nevertheless promoted pre-tRNA maturation in a tRNA-mediated suppression assay. We also uncovered the timing of Trm1-mediated dimethylation with respect to other pre-tRNA processing activities and unexpectedly showed that Sla1 opposes tRNA modification by Trm1. Finally, we provided evidence that both wild-type and catalytically inactive Trm1 are functional in an RNA strand annealing and dissociation assay *in vitro*, suggesting that Trm1 may also act as a tRNA chaperone. These data are thus consistent with the idea that tRNA modification enzymes have retained modification and modification-independent activities throughout evolution and use a combination of these activities to ensure proper tRNA structure and function.

## Results

### Alternate transcriptional start sites yield nuclear– and mitochondrially-targeted Trm1 in *S. pombe*

Trm1 from budding yeast can exist as a nuclear– or mitochondrial-targeted isoform produced from alternate transcription start sites that result in the inclusion or omission of an N-terminal mitochondrial targeting sequence (23). Two alternate transcription start sites have been mapped for fission yeast Trm1: one producing a ∼250 nucleotide 5’ UTR ahead of a more upstream ATG (this isoform will henceforth be referred to as M1, for beginning at the first methionine), and a second with a ∼10 nucleotide 5’UTR ahead of a downstream ATG (which we refer to as M24, beginning at the 24^th^ amino acid) (Figure 1A) (28). While both the longer and shorter isoforms in budding yeast have been shown to target mitochondrially-encoded tRNAs, the N-terminal extension in the longer isoform enhances the efficiency of mitochondrial targeting, resulting in a predominantly mitochondrial-localized protein compared to the nuclear-localized shorter isoform (23). Quantification of the relative abundance of both transcripts in *S. pombe* cells grown in rich media revealed that the shorter transcript giving rise to the M24 isoform was approximately four times as abundant as the mitochondrially-targeted transcript (M1) at steady state levels (Figure 1B). Consistent with the idea that modifications to the tRNA body do not dramatically alter bulk translation, pulse labeling revealed no defects in mitochondrial translation upon Trm1 deletion (Figure S1).

**Figure 1:**
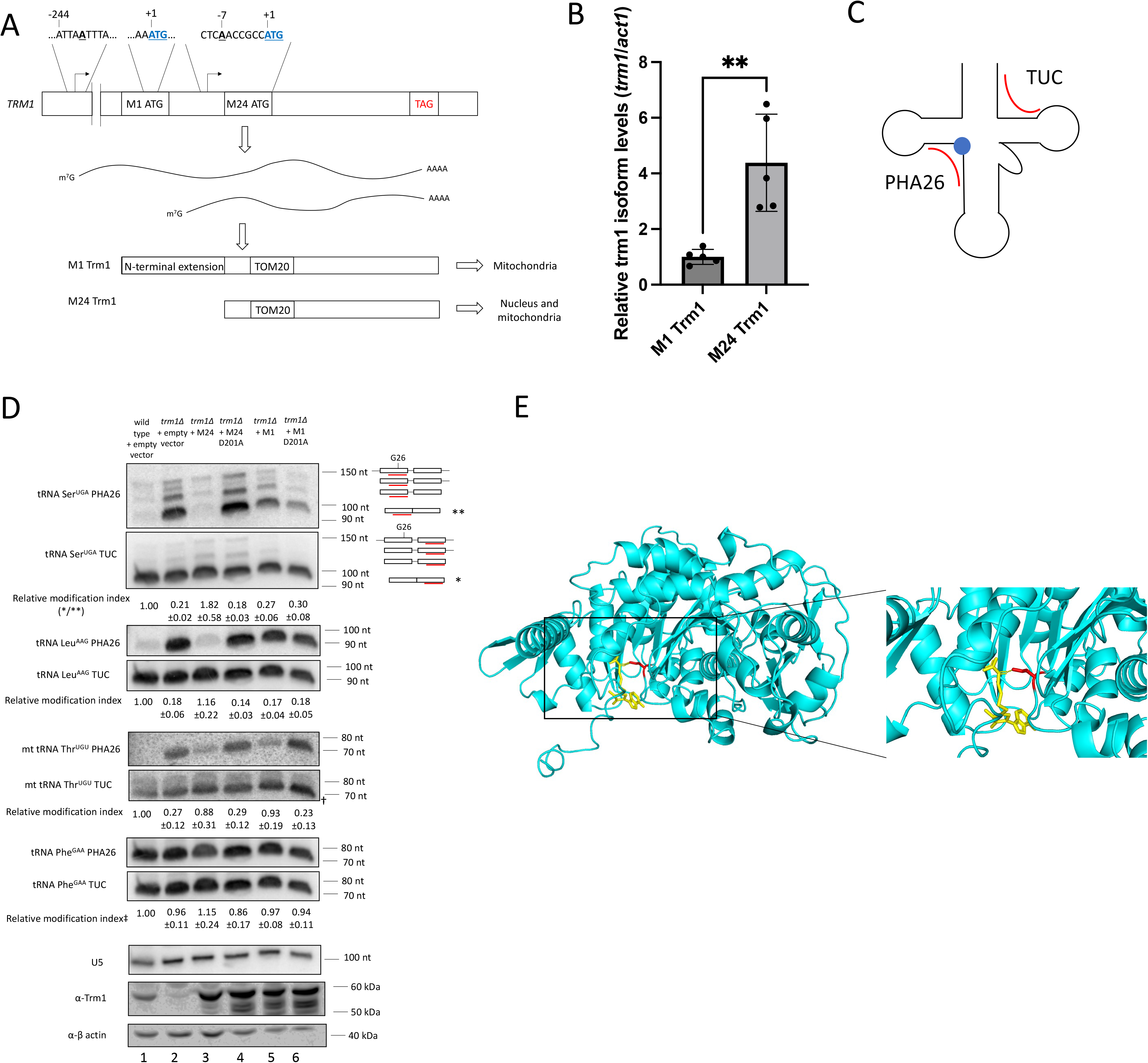
*S. pombe* Trm1 modifies nuclear– and mitochondrial-encoded tRNAs at G26. **A**) Schematic of alternate transcription start sites giving rise to nuclear– and mitochondrial-targeted Trm1 isoforms. The amphiphilic N-terminal sequence in the mitochondrial-targeted isoform enhances the efficiency of mitochondrial targeting, while both isoforms contain a TOM20 binding site for mitochondrial import. B) qRT-PCR of *trm1* mRNA isoforms normalized to *act1* mRNA (mean± standard error, two-tailed unpaired t-test * at p<0.05) (n= 5 biological replicates). C) Schematic depicting binding sites of the PHA26 and TUC probes (red). G26 is represented by a blue circle. D) PHA26 northern blot of nuclear– and mitochondrial-encoded tRNA. Northern blots were stripped and re-probed for U5 as a loading control. Relative modification index represents the TUC signal divided by the PHA26 signal and normalized to a wild type strain (mean±SEM, n= 3 biological replicates). The relative modification index for tRNA Ser^UGA^ was calculated using the signal corresponding to the mature tRNA (**/*). The prominent band corresponding to mt tRNA Thr^UGU^ is indicated (†) to differentiate it from a non-specific background band. ‡Note that although tRNA Phe^GAA^ is not modified, a relative modification index is still provided to demonstrate that the ratio of TUC/PHA26 signal remains unchanged based on the presence of Trm1. Bottom panels: western blot of Trm1 and β-actin in a wild type and *trm1Δ* strain transformed with the indicated plasmids. E) AlphaFold (30) structure prediction of Trm1 aligned to SAH-bound Trm1 from *Pyrococcus horikoshii* (PDB 2EJU) (31). Inset: Hydrogen bonding between SAH (yellow) and D201 (red).

To monitor Trm1-catalyzed modification of nuclear– and mitochondrially-encoded tRNAs, we used an established northern blotting based assay: positive hybridization in the absence of modification at G26 (PHA26) (21, 29), in which dimethylation at G26 impairs hybridization of a probe designed to anneal to the region overlapping G26. A second probe targeting the 3’ end of the tRNA and extending into the TUC stem, which is free of modifications that interfere with probe hybridization, served as an internal normalization for tRNA abundance (Figure 1C). These northern blots supported the expected subcellular targeting of the Trm1 isoforms: M24 Trm1 robustly modified nuclear– and mitochondrial-encoded tRNAs, consistent with its proposed localization to the nucleus and mitochondria, while M1 Trm1 only modified mitochondrially-encoded tRNAs, suggesting that it is solely present in the mitochondria (Figure 1D, lane 5). The ability of overexpressed M24 to rescue mitochondrial tRNA modification levels to a similar degree as overexpressed M1 may result from the increased expression of plasmid-encoded Trm1 relative to endogenous Trm1 (Figure 1D, lane 3), leading to an increase in the amount of Trm1 capable of entering the mitochondria.

These results also confirmed the previously demonstrated substrate specificity of Trm1, where not all G26-containing tRNAs are Trm1 substrates (21). Certain nuclear-encoded tRNAs, including tRNA Ser^UGA^ and tRNA Leu^AAG^ were robustly modified by endogenous Trm1 (Figure 1D, “wild type”, lane 1) and overexpressed M24 Trm1 (Figure 1D, lane 3), while the PHA26 probe annealed to the G26-containing tRNA Phe^GAA^ in a manner that remained unchanged based on the presence of Trm1, consistent with a lack of modification (see figure caption for additional details on quantification of relative modification levels).

### D201 is a key catalytic residue for N2, N2-dimethylation of nuclear– and mitochondrially-encoded tRNAs

To investigate potential modification-independent functions of Trm1, we aligned an AlphaFold (30) prediction of *S. pombe* Trm1 to the structure of an archaeal Trm1 homolog bound to S-Adenosyl Homocysteine (SAH) (31), mutated a conserved aspartic acid directly in the predicted catalytic site to alanine (D201A) (Figure 1E), and again monitored Trm1-catalyzed modification of nuclear– and mitochondrially-encoded tRNAs by the PHA26 assay. As expected from the proposed role of this conserved aspartic acid in deprotonating G26 for nucleophilic attack of SAM (31), the D201A mutant showed the same lack of modification of nuclear– and mitochondrial-encoded tRNAs as *trm1Δ* cells transformed with an empty vector, despite similar levels of protein accumulation as the wild type overexpressed isoform (Figure 1D).

To rule out defects in tRNA binding contributing to a lack of modification by D201A, we purified recombinant wild type and D201A M24 Trm1 and measured *in vitro* binding affinity to pre-tRNA Ser^UGA^ using electrophoretic mobility shift assays (EMSA). Wild type and D201A exhibited comparable binding affinity, suggesting that disruption of the putative catalytic site does not significantly impair tRNA binding (Figure 2A, B, Figure S2). We note that the supershift binding patterns (defined as complexes migrating at a higher molecular weight than the initial binding event indicated with an asterisk), were slightly different between wild type Trm1 and D201A. We anticipate that the initial binding event represents the physiologically relevant binding event, while the additional bound conformations at higher concentrations may result from more than one protein bound per RNA at sites that are not the primary binding site. The presence of multiple supershift bands with the D201A mutant may be therefore due to unanticipated differences in substrate accommodation occurring upon non-physiologically relevant binding events when one tRNA is bound by multiple proteins.

**Figure 2:**
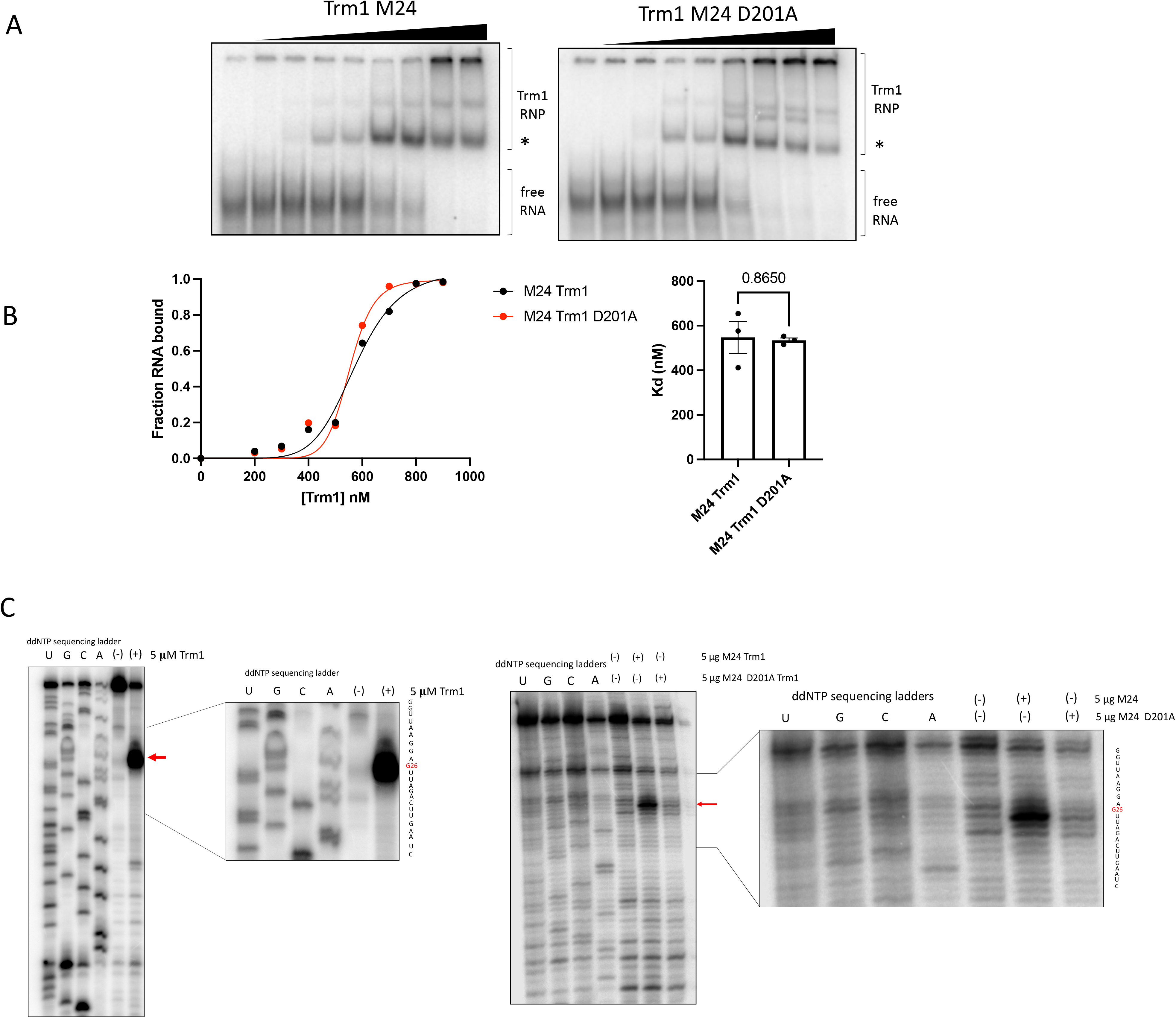
D201A supports *in vitro* tRNA binding, but not methylation. **A**) Representative EMSAs of Trm1 M24 and M24 D201A with radiolabeled pre-tRNA Ser^UGA^. The asterisk represents the initial binding event. B) Representative binding curves (left) and *K*_D_ values of A) (right) (mean±SEM, two-tailed unpaired *t* test, n= 3 technical replicates) (see Figure S2 for all binding curves). C) Primer extension of *in vitro* methylated tRNA Ser^UGA^ with 5 μM (left) or 5 μg (right) recombinant Trm1. The RNA sequences flanking G26 are indicated and the modification-dependent RT stop is indicated by a red arrow.

We also set up *in vitro* methylation reactions with pre-tRNA Ser^UGA^, recombinant Trm1, and S-Adenosyl methionine (SAM), and measured modification efficiency by primer extension, as dimethylation on the Watson-Crick face is sufficient to cause a reverse transcriptase (RT) stop (18, 32). The lack of modification by D201A *in vitro* (Figure 2C), evident by the lack of RT stop, and by PHA26 northern blotting *in vivo* (Figure 1D) confirmed that D201A is indeed catalytically inactive, thus enabling further studies into potential catalytic-independent functions of Trm1.

### Trm1 promotes tRNA-mediated suppression through catalytic and catalytic-independent activities

tRNA-mediated suppression is an established system in *S. pombe* that has been used to test various aspects of pre-tRNA maturation including 5’ leader, 3’ trailer and intron processing and tRNA modification (reviewed in (33, 34)). The assay relies on a mutation to the anticodon of tRNA Ser^UCA^, allowing stop codon readthrough of a nonsense mutation in the AIR carboxylase gene which, when fully functional, prevents the accumulation of a red metabolic intermediate (33, 35). The G35C mutation that enables nonsense decoding also impairs anticodon-intron base pairing in the suppressor pre-tRNA, resulting in a misfold that increases susceptibility of the pre-tRNA to nuclear pre-tRNA quality control and exosome-mediated decay (16). Thus, this assay has been used previously to monitor pre-tRNA susceptibility to nuclear surveillance (16). Suppressor pre-tRNA degradation linked to mutations predicted to cause misfolding can be rescued by overexpression of pre-tRNA processing factors, enabling identification and insight into the mode of action of pre-tRNA binding proteins, including the RNA chaperone La (15, 16). We and others have previously demonstrated that Trm1 is active in promoting tRNA-mediated suppression (18, 21), although this could be due to a stabilizing effect from the modification, a separate pre-tRNA maturation or stabilizing activity that functions independently from modification, or a combination of these.

We found that both wild type and catalytically inactive nuclear-targeted (M24) Trm1 promoted suppression of the tRNA Ser^UCA^ allele in a *sla1Δ* background, suggesting that modification is not strictly required for suppression activity and that Trm1 promotes pre-tRNA maturation even in the absence of catalysis (Figure 3A, B). Addition of the U47:6C (yeast strain ySH18) or C40U, U47.3C, and C47.6U (yeast strain yYH1) mutations, which are associated with increased reliance on the RNA chaperone activity of La (17, 36), resulted in partial tRNA-mediated suppression by wild type Trm1 but not D201A (Figure 3B, C). These data are suggestive of two activities for Trm1: its established methyltransferase activity, which has been previously shown to stabilize tRNA structure (18), and a modification-independent activity. The catalytic-independent function is sufficient for suppression activity in the context of the tRNA Ser^UCA^ allele, whereas both activities are required for more defective suppressor tRNA alleles.

**Figure 3:**
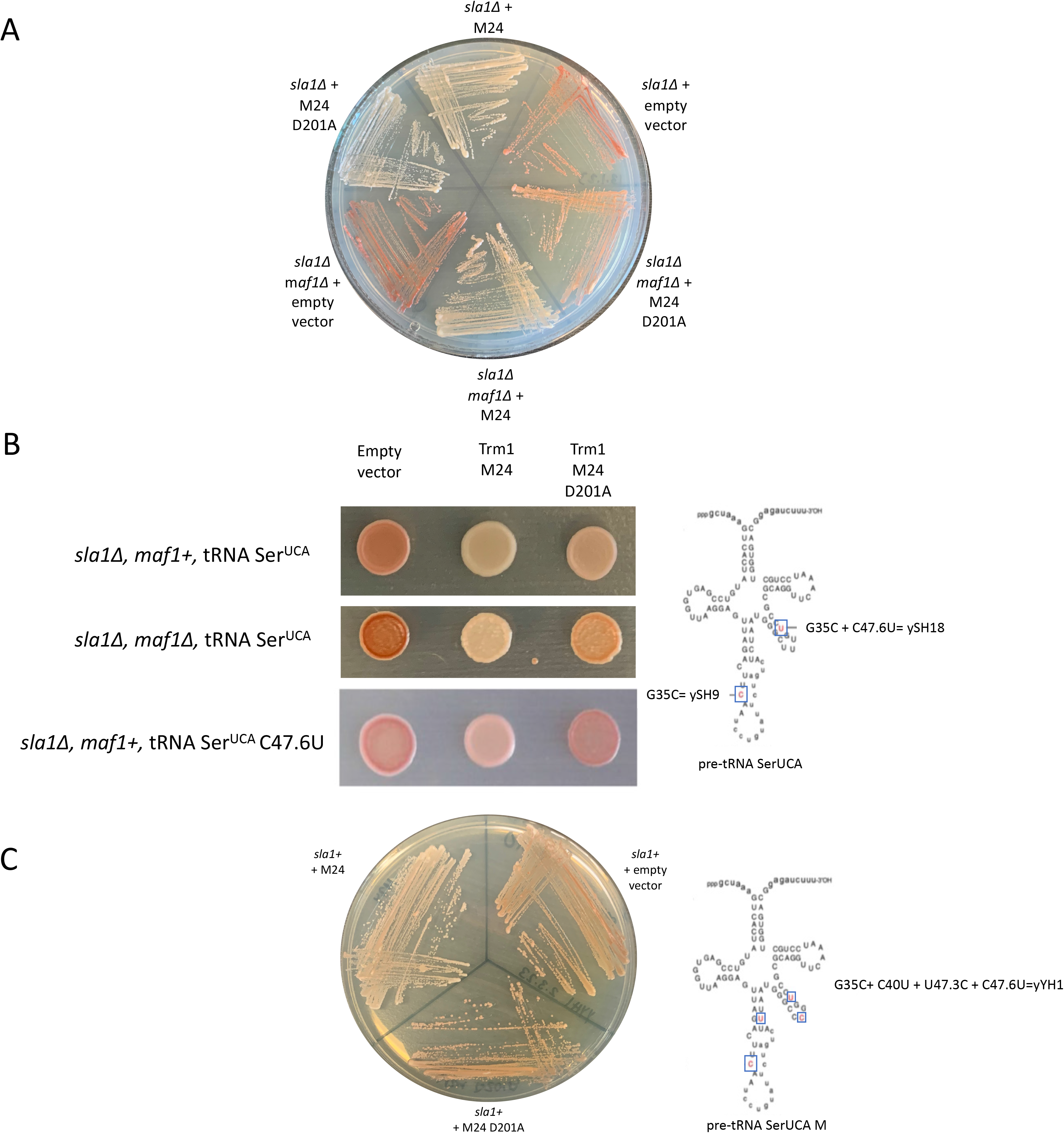
Trm1 promotes tRNA-mediated suppression independent of catalytic activity. **A**) tRNA-mediated suppression with wild type and catalytically inactive Trm1 in a *sla1Δ* and *sla1Δ maf1Δ* background. B) tRNA-mediated suppression with wild type and catalytically inactive Trm1 in a *sla1Δ* and *sla1Δ maf1Δ* background with increasingly defective suppressor tRNA alleles. Right: schematic of the G35C (yeast strain ySH9) and G35C C47.6U (yeast strain ySH18) suppressor tRNA alleles. C) tRNA-mediated suppression of the MSer suppressor tRNA allele with wild type and catalytically inactive Trm1 in a *sla1+* strain. Right: Schematic of the MSer suppressor tRNA allele integrated into the yeast strain yYH1.

It has also been reported that deletion of the RNA Polymerase III repressor Maf1 causes anti-suppression in the tRNA mediated suppression assay, which at first was noted to be counter-intuitive with the increase in tRNA transcription observed upon Maf1 deletion (21, 37). However, this anti-suppression phenotype can be explained by hypomodification of the suppressor tRNA by Trm1, as Trm1—and presumably other pre-tRNA binding proteins— become limiting with the resultant increase in the pre-tRNA pool (21). We found that suppression of tRNA Ser^UCA^ is achieved by wild type and catalytically inactive Trm1 in the *maf1Δ* strain, although we note that the degree of suppression, as determined by a lack of red pigment accumulation, was greater for wild type Trm1 compared to the catalytically inactive mutant (Figure 3A, B). In the model whereby Trm1 becomes limiting in the absence of Maf1, suppression by D201A suggests that it is not only the Trm1-catalyzed modification that becomes limiting, but also a catalytic-independent function for Trm1. Still, the additional suppression achieved by wild type Trm1 compared to the catalytically inactive mutant supports a stronger role for the Trm1-catalyzed modification than a secondary catalytic-independent activity.

To further investigate how Trm1 promotes tRNA-mediated suppression, we examined suppressor tRNA processing and modification by northern blotting (Figure 4A). Probes against the tRNA Ser^UCA^ intron were used to detect the precursor, which migrated as 2 bands in the absence of the yeast La protein Sla1: a leader-containing species that migrated as a higher molecular weight smear due to 3’ exonucleolytic nibbling (represented by * for pre-tRNA Ser^UGA^ and pre-tRNA Lys^CUU^), and a leader– and trailer-processed species (**) (Figure 4A). In contrast, the presence of Sla1 resulted in 3 visible pre-tRNA processing intermediates corresponding to the nascent leader– and trailer-containing pre-tRNA, the leader-processed and trailer-containing pre-tRNA, and the fully end processed pre-tRNA (Figure 4B, pre-tRNA Lys^CUU^). We also used a probe overlapping the exon junction to detect the mature suppressor tRNA. As the probe targeting the exon junctions overlaps the anticodon loop (ACL), which contains post-transcriptional modifications such as m^3^C32, i^6^A37, and t^6^A37 which are known to influence probe binding (29, 38, 39), we designed a probe targeting the ACL of wild type tRNA Ser^UGA^ to test whether the presence of Trm1 impacts certain ACL modifications (Figure S3). We did not observe any differences in probe binding due to the presence or absence of endogenous or overexpressed Trm1, suggesting that Trm1 does not influence ACL modifications that are sensitive to northern blotting.

**Figure 4:**
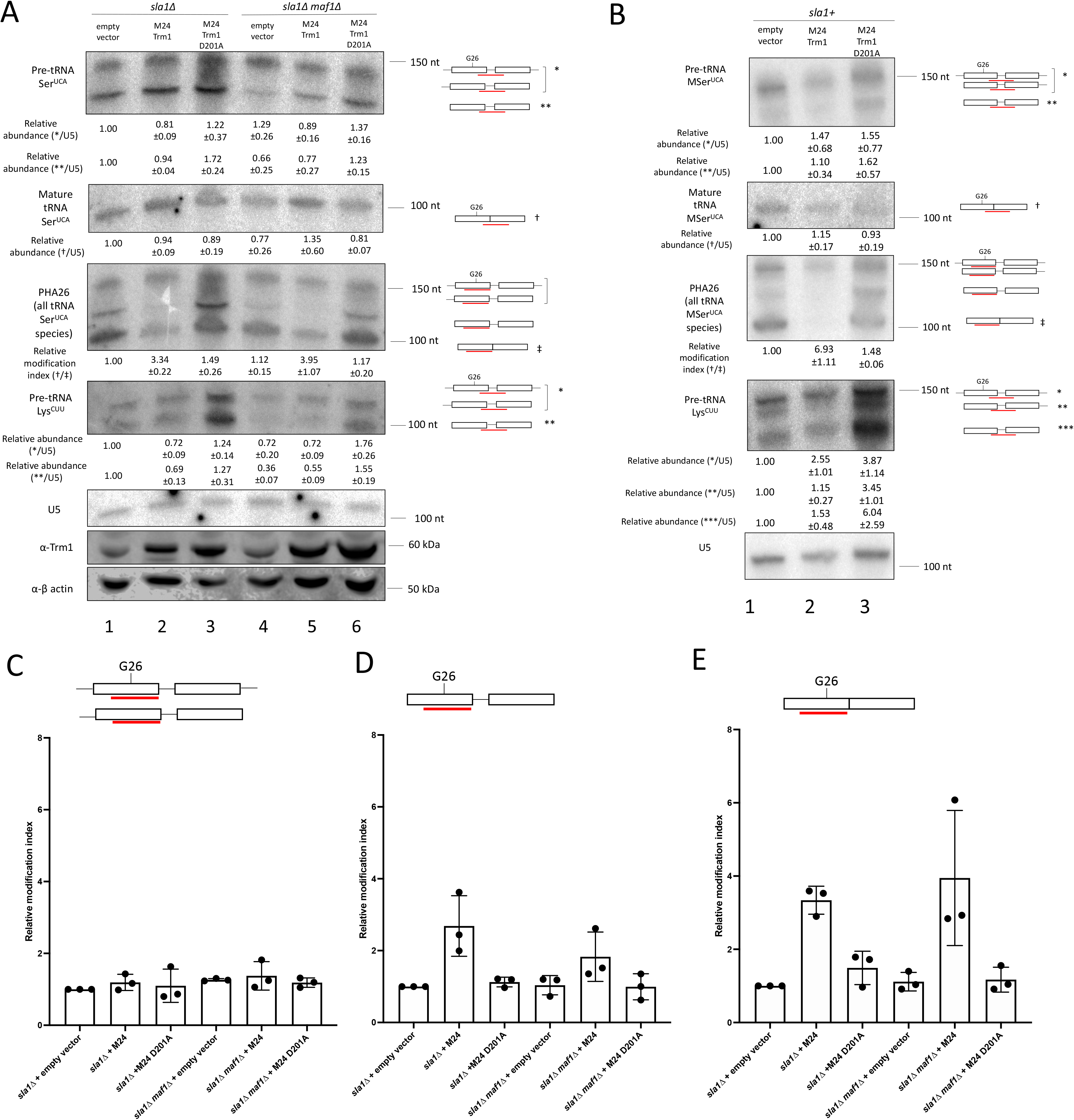
Trm1 modifies end-processed, intron-containing pre-tRNAs. **A**) Northern blot of tRNA Ser^UCA^ processing intermediates and G26 modification status and 3’ end protection of pre-tRNA Lys^CUU^ and western blot of Trm1 and β-actin in a *sla1Δ* and *sla1Δ maf1Δ* strain transformed with the indicated plasmids. * refers to the nascent and 3’ end-processed suppressor pre-tRNA, ** is the intron-containing, end-processed suppressor pre-tRNA, † is the probe overlapping the splice junction, and ‡ is the PHA26 probe overlapping G26 (mean±SEM, n= 3 biological replicates). B) Northern blot of tRNA Ser^UCA^ processing intermediates and G26 modification status and 3’ end protection of pre-tRNA Lys^CUU^ in a *sla1+* strain transformed with the indicated plasmids (mean±SEM, n= 3 biological replicates). C-E) Quantification of relative modification at G26 in A), relative to the *sla1Δ* strain expressing an empty vector. Relative modification was calculated by normalizing the PHA26 signal (overlapping G26) to probes targeting the intron-containing precursor (C and D) and mature suppressor (E) (mean±SEM, n= 3 biological replicates).

tRNA-mediated suppression of tRNA Ser^UCA^ by human or *S. pombe* La is largely due to 3’ trailer binding and protection of the suppressor pre-tRNA from exosome-mediated decay, which is characterized by the stabilization of a 3’ trailer-containing suppressor pre-tRNA species with ectopic La expression in a *sla1Δ* strain relative to an empty vector (15, 16) (also see Figures 5 and 6). 3’ end protection by La and La-related Proteins (LARPs) is typically monitored by looking for stabilization of the 3’ trailer-containing pre-tRNA Lys^CUU^, due to the increased resolution of pre-tRNA processing intermediates relative to pre-tRNA Ser^UGA^ (16, 40, 41). tRNA Lys^CUU^ is also modified by Trm1, making it an ideal target to test whether Trm1 functions in tRNA-mediated suppression through 3’ end protection (21). As might be expected based on previous work linking recognition of tRNAs by Trm1 to the D-loop (42), we observed no differences in 3’ end protection upon addition of wild type Trm1, as evident by the lack of stabilization of the 3’ trailer-containing species relative to the empty vector (Figure 4A, pre-tRNA Ser^UCA^ and endogenous pre-tRNA Lys^CUU^, compare to stabilization of the nascent pre-tRNA with Sla1, see below). This suggests that unlike the La protein, wild type Trm1 promotes tRNA-mediated suppression in the absence of 3’ end protection, much like what has been described for tRNA-mediated suppression by La-related Proteins (41). However, we noted that stabilization of the 5’ and 3’ processed pre-tRNA species occurs upon overexpression of D201A, which supports the idea that in the absence of catalysis, Trm1 may remain bound to and stabilize the pre-tRNA (Figure 4A, lanes 3 and 6; Figure 4B, lane 3). The accumulation of pre-tRNA following overexpression is also consistent with our *in vitro* results suggesting that the D201A mutation does not significantly impair Trm1’s tRNA binding activity (Figure 2A, B, S2), as a loss of high affinity tRNA binding would not be expected to manifest as a pre-tRNA accumulation phenotype, although we note that we have not investigated how other factors involved in pre-tRNA processing might be directly or indirectly affected by D201A overexpression.

**Figure 5:**
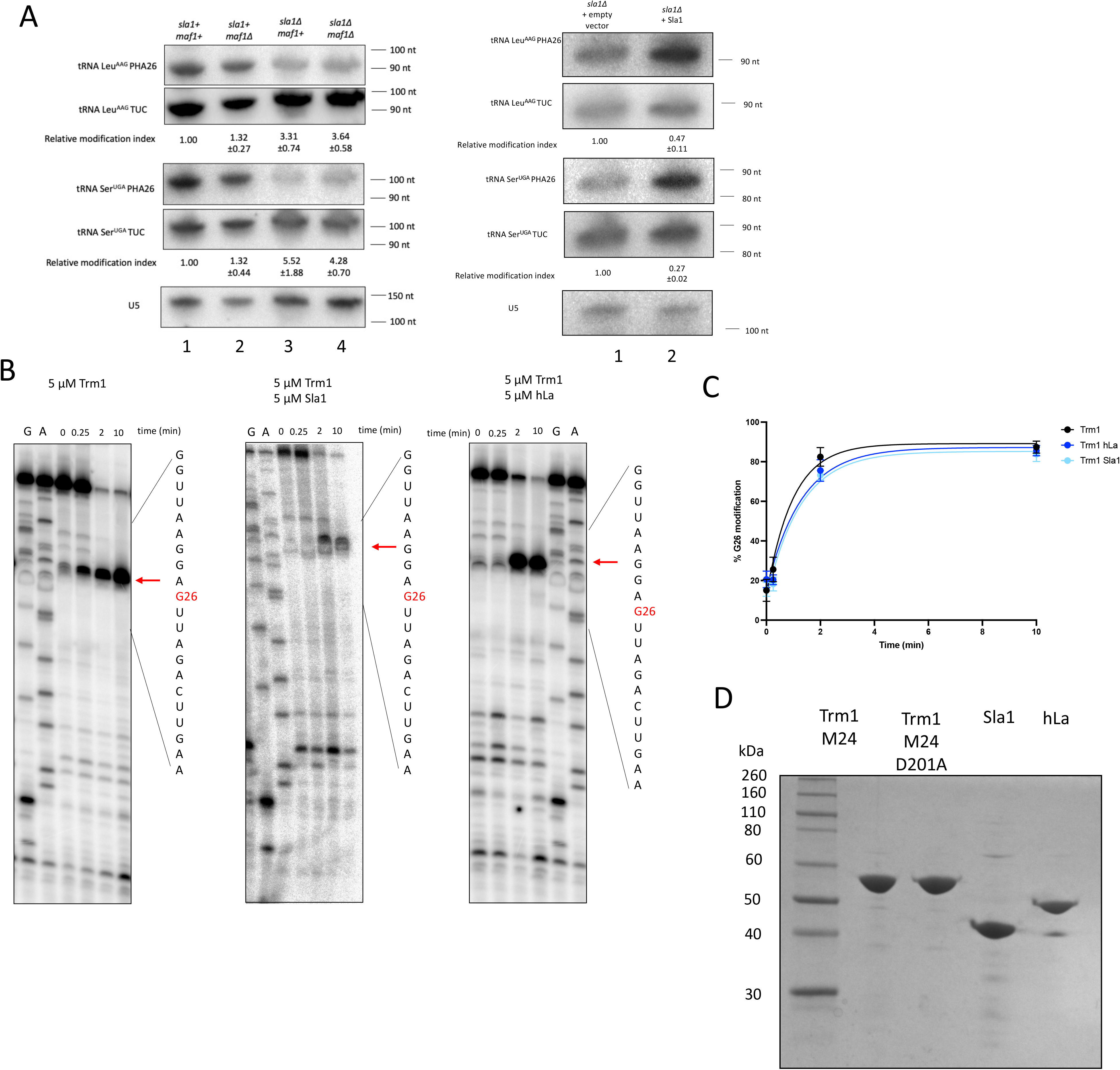
La and Trm1 influence pre-tRNA end-processing and G26 dimethylation. **A**) PHA26 northern blots of nuclear-encoded tRNA in wild type, *maf1Δ, sla1Δ,* and *sla1Δ maf1Δ* cells (left) and *sla1Δ* strains transformed with an empty vector or Sla1 (right). Northern blots were stripped and reprobed for U5 as a loading control. Relative modification index represents the TUC signal divided by the PHA26 signal and normalized to a wild type or empty vector-expressing strain (mean±SEM, n= 3 biological replicates) B) Primer extension of *in vitro* methylated tRNA Ser^UGA^ pre-incubated with no protein, *S. pombe* La (Sla1), or human La (hLa) prior to incubation with Trm1. Red arrows indicate the reverse transcriptase stop corresponding to G26. C) Quantification of *in vitro* methylation of tRNA Ser^UGA^ over time, calculated as the ratio of the G26 RT stop to full length tRNA (mean±SEM, n= 4 independent replicates). Points were fit to a non-linear specific binding curve (One-phase association with least squares fit). D) SDS-PAGE analysis of recombinant proteins used in this study. 1 μg of each protein was assessed for quantity and purity and visualized with Coomassie blue stain.

**Figure 6:**
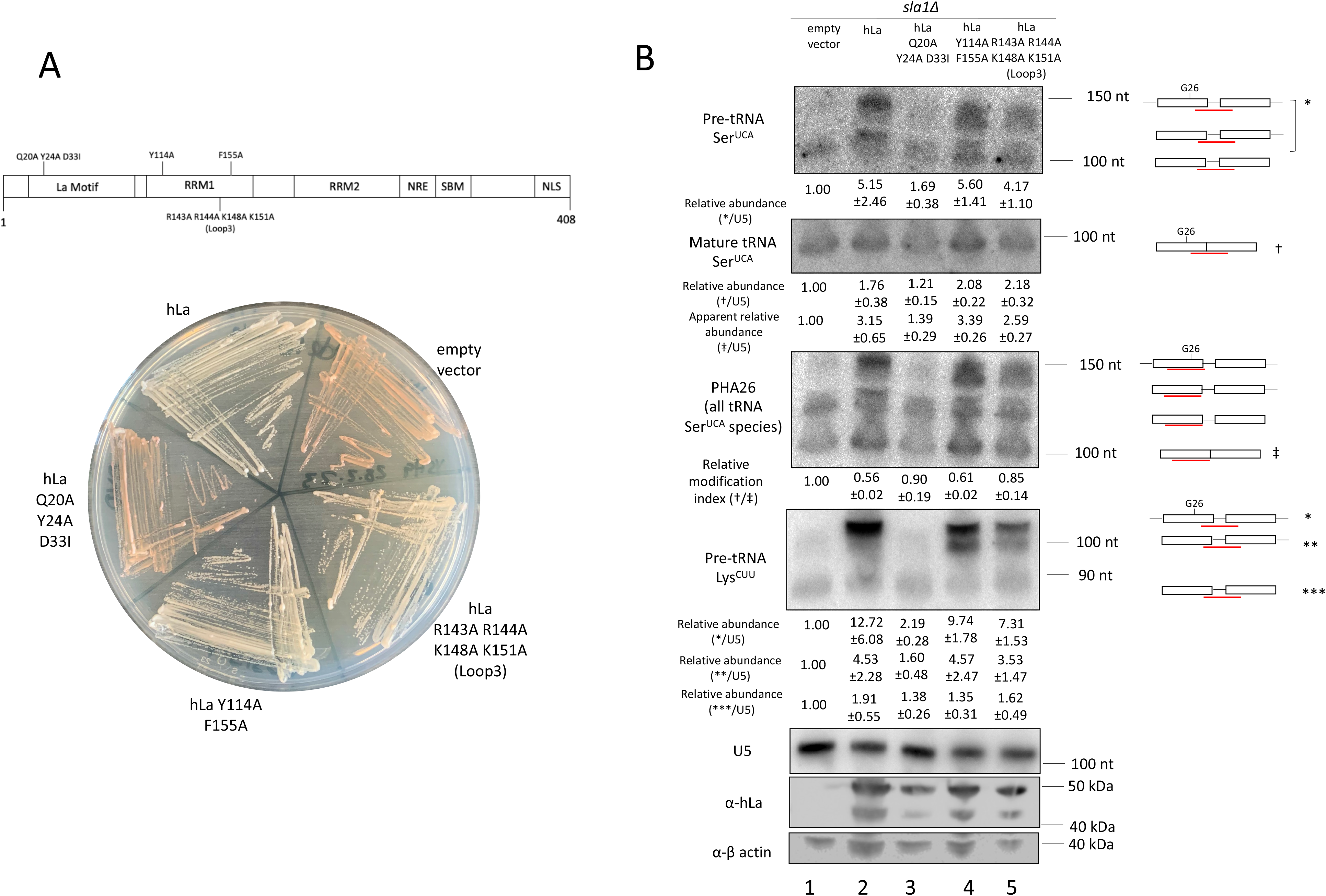
Crosstalk between La and Trm1 is linked to La-pre-tRNA binding modes. **A**) Schematic of human La (hLa) domains with indicated mutations and tRNA-mediated suppression of wild type and mutant hLa constructs in a *sla1Δ* background. B) Northern blot of tRNA Ser^UCA^ processing intermediates and G26 modification status and 3’ end protection of pre-tRNA Lys^CUU^. Apparent relative abundance refers to intensity of the PHA26 probe normalized to U5. A western blot for human La and β-actin is provided below. * refers to the nascent leader– and trailer-containing suppressor pre-tRNA, † is the TUC probe overlapping the splice junction, and ‡ is the PHA26 probe overlapping G26. Apparent relative abundance was calculated by dividing signal intensity from the PHA26 probe targeting the mature suppressor tRNA to signal intensity of U5, while relative modification index was calculated by dividing signal intensity of the TUC probe targeting the mature suppressor tRNA divided by the PHA26 probe targeting the mature suppressor tRNA (mean±SEM, n= 3 biological replicates).

We also performed the PHA26 assay to investigate the timing of Trm1 modification in tRNA-mediated suppression (Figure 4A, B, third panel). We observed that overexpression of Trm1 in the *trm1+* suppressor-tRNA containing strain resulted in a decrease in signal intensity with the PHA26 probe, consistent with previous reports that endogenous Trm1 levels are insufficient for full tRNA modification (21). However, overexpression of the catalytically inactive D201A mutant resulted in comparable modification levels in this strain to an empty vector control (Figure 4A, lanes 3 and 6, Figure 4B, lane 3). The PHA26 assay also provided information on the timing of Trm1 modification with respect to pre-tRNA processing activities: we observed no change in PHA signal intensity upon Trm1 expression in the trailer-processed, leader-containing pre-tRNA (Figure 4A, top band in third panel), suggesting that it is not a substrate for modification, but noted a decrease in signal intensity for the leader– and trailer-processed pre-tRNA and mature tRNA, supporting a previously proposed model that Trm1 modification occurs after pre-tRNA end processing (18) (Figure 4C-E). Still, although Trm1 modification starts to occur at the pre-tRNA level, tRNAs were not fully modified (which we infer from the greater decrease in PHA26 signal intensity between the empty vector and overexpressed Trm1) until the mature tRNA stage. We noted the same pattern, with modification evident on the end-processed, intron-containing pre-tRNA but not the nascent pre-tRNA, for the endogenous tRNA Ser^UGA^ in a wild type and *trm1Δ* strain (Figure 1D).

As Sla1 promotes 5’ leader processing prior to 3’ trailer processing, resulting in the accumulation of a 3’ trailer-containing pre-tRNA species (14), we repeated these experiments in a wild type (*sla1+*) strain to uncouple Sla1-dependent changes in tRNA 3’ end processing from Trm1-mediated stabilization. For these experiments, we used the MSer suppressor tRNA allele (Figure 3C) (40), which has more mutations predicted to cause misfolds than the G35C and U47:6C alleles and for which endogenous Sla1 levels are insufficient for tRNA-mediated suppression (40). In this *sla+* background only wild type Trm1 was capable of suppressing MSer, much like the U47:6C allele (Figure 3C), supporting the importance of G26 dimethylation in imparting additional tRNA stability for more substantially misfolded tRNAs. Aside from increased accumulation of 3’ trailer-containing pre-tRNA species (Figure 4B, top 2 bands of pre-tRNA Lys^CUU^), we observed similar patterns to the *sla1Δ* strain: suppression by wild type Trm1 was not due to increased 3’ end protection and Trm1 modification occured on end-processed, intron containing pre-tRNAs (Figure 4B, PHA26). These data support the idea that Trm1-catalyzed modification and activity in the tRNA-mediated suppression assay are not affected by the order of 3’ end processing, as it remains unchanged based on the presence of Sla1.

### The La protein decreases Trm1-catalyzed modification

While previous reports indicate that tRNAs become hypomodified at G26 upon Maf1 deletion (21), we did not observe any modification differences with and without endogenous Maf1 (Figure 4A,C-E). Since those previous measurements were performed in a *sla1+* strain and our data are from a *sla1Δ* strain, we examined the relationship between Sla1 and Trm1-dependent modification. Strikingly, we detected a substantial increase in G26 modification upon Sla1 deletion for leucine and serine tRNAs, suggesting a possible crosstalk between Sla1 and Trm1 for pre-tRNA binding (Figure 5A). Importantly, re-expression of Sla1 in a *sla1Δ* strain was sufficient to decrease G26 modification (Figure 5A, right panel), while overexpression of wild type Trm1 in a *sla1+* strain increased G26 modification, confirming previous results that endogenous Trm1 is insufficient to fully modify the cellular pool of tRNAs (21) (Figure 4B PHA26 panel, Figure S4). Conversely, overexpression of Sla1 in a *sla1+* strain resulted in no changes in G26 modification, possibly because endogenous Sla1 levels already outcompete endogenous Trm1 levels (43, 44), with increased Sla1 expression exerting negligible effects on tRNA processing and modification (Figure S5). Notably, the degree to which G26 modification increased upon Trm1 overexpression was greater in a *sla1+* strain than a *sla1Δ* strain, suggesting that a larger fraction of tRNAs are already fully modified in the absence of Sla1 (Figure S4).

To investigate whether La binding shields pre-tRNAs from Trm1 modification of nuclear-encoded pre-tRNAs, we pre-incubated pre-tRNA with and without recombinant Sla1 or human La (hLa) and set up an *in vitro* methylation time course to examine changes in G26 dimethylation. We observed slightly decreased methylation in the presence of Sla1 and hLa, particularly at earlier time points in the reaction (Figure 5B, C, compare timepoint t=0.25 minutes between gels), consistent with our *in vivo* data demonstrating that La decreases Trm1 modification. We note that further kinetic analyses must be conducted to determine differences in the rate of modification with and without La. As the methylation reaction occurs in the absence of any cellular factors that promote tRNA processing, including other modification enzymes and end-processing endo and exonucleases, this is suggestive of La itself acting as a barrier to Trm1-mediated methylation, rather than decreasing dimethylation indirectly, through its roles in facilitating pre-tRNA processing.

As the La protein has been reported to make multiple contacts to pre-tRNAs—engagement of the uridylate trailer and pre-tRNA body (15, 16, 45)—we took advantage of previously characterized mutants of hLa to map inhibition of tRNA modification to La’s various domains and RNA binding modes (Figure 6A). We measured tRNA-mediated suppression activity, steady-state precursor and mature tRNA levels, and G26 modification status for the suppressor tRNA Ser^UCA^ in a *sla1Δ* strain transformed with wild type or mutant hLa (Figure 6). As has been previously reported, mutations to uridylate-binding residues in the La motif (Q20A Y24A D33I) led to decreased suppression activity. In contrast, mutations to the RRM1 (Y114A F155A and RRM1 loop-3 R142A R143A K148A K151A), which result in impaired RNA chaperone activity but unchanged uridylate binding (15), exhibited no defects in the tRNA-mediated suppression assay, in agreement with the idea that tRNA Ser^UCA^ predominantly requires 3’ end protection, but not RNA chaperone activity, for complete suppression (16) (Figure 6A). This was evident from stabilization of the top suppressor tRNA band on the northern blots, corresponding to the nascent, 5’ leader– and 3’ trailer-containing pre-tRNA species, for wild type and RRM1 hLa mutants (Figure 6B).

We observed a robust decrease in G26 modification of the suppressor tRNA in strains transformed with wild type hLa and the RRM1 mutant Y114A F155A, which displays decreased RNA chaperone activity but has no pre-tRNA binding defects (16). On the other hand, we observed no differences in modification between the empty vector and the uridylate-binding mutant, and a modest decrease in modification compared to the empty vector by the RRM1 loop-3 R142A R143A K148A K151A mutant, which is defective in RNA chaperone activity and has impaired but not ablated pre-tRNA binding that is not attributed to La’s uridylate binding mode (15, 17) (Figure 6B). These results suggest that crosstalk between La and Trm1 modification correlates with La’s pre-tRNA binding mode, with mutations that disrupt uridylate or tRNA body binding, which contribute additively to pre-tRNA affinity (15), leading to increased modification at G26. Notably, these results also revealed that northern blot-based measurements of hLa-dependent increases in mature suppressor tRNA levels could arise from a combination of changes in mature tRNA levels and differences in G26 modification, for tRNA probes that overlap with this site. Consistent with this, changes in apparent mature suppressor tRNA levels in the presence or absence of wild type or mutant hLa were less pronounced when using a probe that does not overlap with G26 (Figure 6B, compare relative mature tRNA Ser^UCA^ to apparent relative mature tRNA Ser^UCA^). Thus, mature suppressor tRNA levels are not the sole determinant that can be used to predict or assign meaning to tRNA-mediated suppression activity, providing additional evidence for the hypothesis that an increase in specific tRNA activity due to modification or folding is linked to tRNA-mediated suppression activity, rather than suppression due solely to suppressor tRNA accumulation.

### Trm1 promotes RNA strand annealing and dissociation *in vitro*

As our data point towards Trm1 possessing a catalytic-independent function that promotes tRNA functionality *in vivo*, we considered the possibility that Trm1 might function as an RNA chaperone, much like the bacterial tRNA modifying enzymes TruB and TrmA (25, 26). We employed a previously established, *in vitro* FRET-based RNA annealing and dissociation assay that has been used to characterize RNA chaperone activity for La, La-related proteins and other established RNA chaperones such as Hfq and StpA (17, 41, 46–48). This assay uses complementary RNA oligos 5’ end labeled with Cy3 or Cy5, such that annealing of the oligos results in FRET between the two fluorophores, from which a rate of annealing is calculated (*k_ann_*), followed by the addition of excess unlabeled competitor to assess strand dissociation activity (*k_SD_*), which is accompanied by a decrease in FRET between the two fluorophores (Figure 7A). In the absence of strand dissociation, as measured by an increase in FRET following competitor addition, the rate constant calculated in the second phase is considered *k_ann2_*. Consistent with previous data (17, 41), strand annealing and strand dissociation were enhanced by the addition of recombinant human La (Figure 7B, C). We also observed strand annealing and dissociation for both wild type and catalytically inactive Trm1 (Figure 7B, C). Our *in vivo* data demonstrating that Trm1 is active in a tRNA-mediated suppression assay independent of catalysis, coupled with our *in vitro* data supporting an RNA chaperone-like activity, suggest that Trm1 may influence pre-tRNA maturation through a combination of structure-stabilizing modifications and protein-mediated RNA annealing and unwinding, similar to prokaryotic tRNA modification enzymes (25, 26).

**Figure 7:**
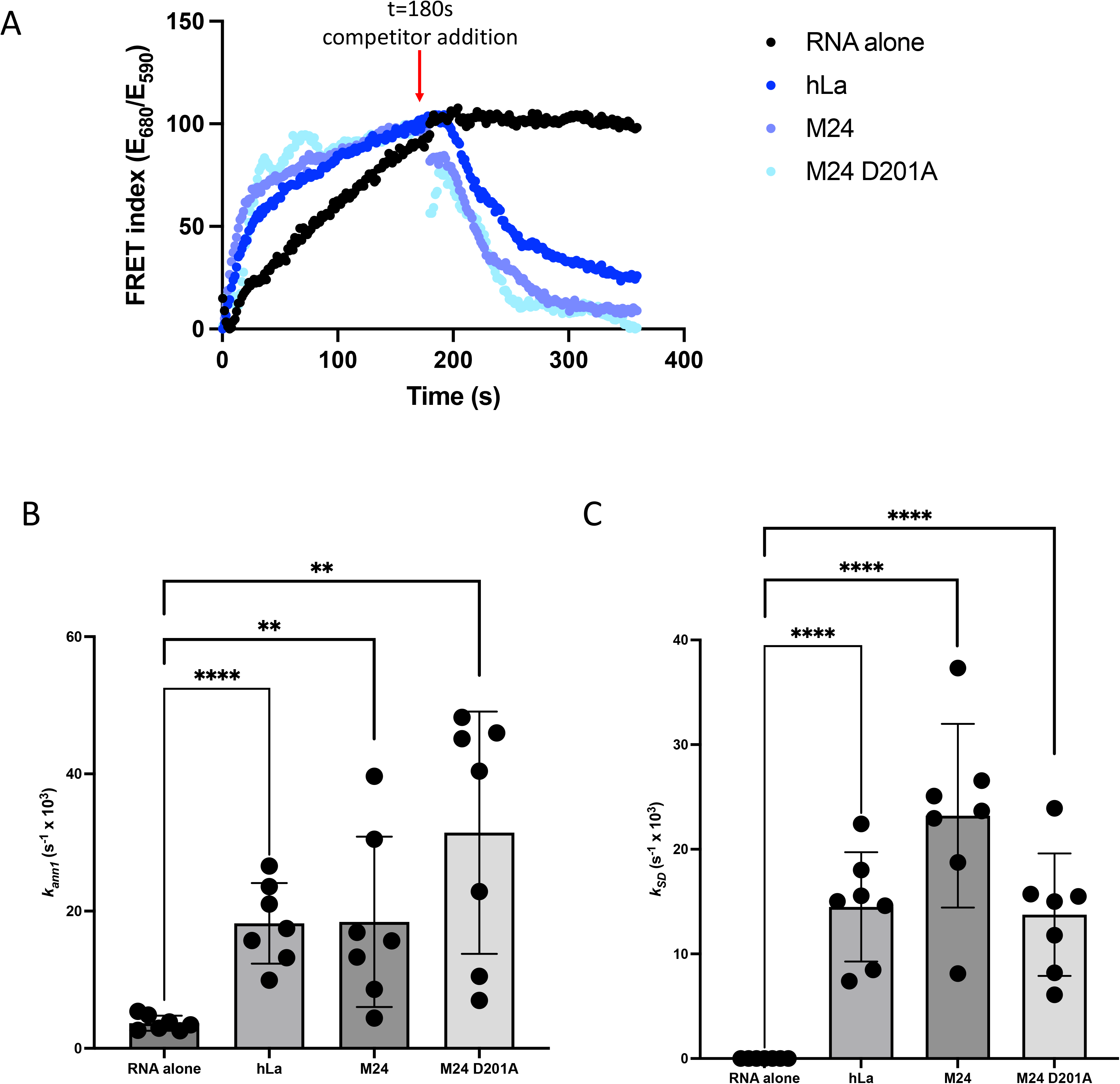
Trm1 exhibits RNA strand annealing and dissociation *in vitro*. **A**) RNA strand annealing (phase I) and dissociation (phase II), as indicated by changes in FRET index (emission at 680 nm/emission at 590 nm) between the Cy3– and Cy5-labeled substrates. Phase II was initiated with the addition of an unlabeled competitor RNA at t=180 seconds. Representative traces are shown. B-C) Strand annealing (B) and dissociation (C) rate constants calculated with RNA alone, recombinant hLa, or wild type or catalytically inactive M24 Trm1 (mean± standard error, two-tailed unpaired t-test * at p<0.05, ** at p<0.01, *** at p<0.001) (n= 7 independent replicates).

## Discussion

In this work, we investigated the interplay between the tRNA modification enzyme Trm1 and the established RNA chaperone La, and provided evidence for a catalytic-independent function of Trm1 with functional overlap with genuine La proteins. Similar to La, we anticipate that an RNA chaperone activity linked to a catalytic-independent function of Trm1 may promote pre-tRNA folding in the nucleus, while Trm1-catalyzed dimethylation is particularly important for additional stabilization to resolve more severe misfolds. This model is consistent with our observations comparing tRNA-mediated suppression of various suppressor tRNA mutants: the G35C suppressor tRNA mutation (Ser^UCA^) only results in a misfold at the pre-tRNA level, due to altered base-pairing with the intron which is later removed, and so the mature suppressor tRNA possesses the same fold as endogenous tRNA Ser^UGA^. In contrast, adding the U47:6C or C40U, U47.3C, and C47.6U mutations lead to a misfold that persists in the mature tRNA in the cytoplasm, compounded by the fact that the degree of misfold in U47:C or C40U, U47.3C, and C47.6U is also greater than that of tRNA Ser^UCA^. Therefore, while the catalytic-independent activity alone is sufficient for Trm1 associated tRNA Ser^UCA^ stabilization and folding in the nucleus, more defective suppressor tRNA alleles may require both Trm1-mediated pre-tRNA stabilization in the nucleus and the Trm1-catalyzed modification that persists on the mature tRNA in the cytoplasm for full suppression activity (Figure 3).

We have previously noted a slight charging defect on the endogenous parent of the tRNA Ser^UCA^ suppressor tRNA (tRNA Ser^UGA^) in the context of Trm1 deletion, as well as altered tRNA structure by *in vitro* chemical probing in the absence of G26 modification (18), consistent with tRNA misfolds linked to mature tRNA function being influenced by G26 dimethylation. Given the overlap in cellular localization and apparent functional and genetic crosstalk with the nuclear La protein, we favor a model in which a Trm1 RNA chaperone function associated with the Trm1 D201A mutant may help protect pre-tRNAs from nuclear surveillance, similar to La. In such a scenario, the act of Trm1 binding pre-tRNAs (in the context of modification or a modification-independent activity) might serve to protect the pre-tRNA from nuclear degradation, in an analogous manner to how La binding to the 3’ uridylate trailer prevents nuclear surveillance or EF1A competes with the rapid tRNA decay machinery in the cytoplasm for access to mature tRNAs to prevent their degradation (16, 49). Still, it remains to be found what proportion of pre-tRNAs benefit from Trm1-catalyzed modification versus the catalytic-independent function, and how Trm1-mediated pre-tRNA stabilization influences downstream tRNA maturation activities to promote tRNA stability and function in translation.

One unexpected result was the apparent inhibition of tRNA modification at G26 by the presence of Sla1, which we validated *in vitro* showing that Sla1 and hLa restrict access of Trm1 to a tRNA substrate. Previously, Trm1 and La were hypothesized to function redundantly in the tRNA biogenesis pathway (18, 19), with La-mediated pre-tRNA stabilization thought to increase the window in which Trm1 could modify pre-tRNAs in the nucleus (18). While it is indeed likely that Trm1 and La have redundant functions — both promote pre-tRNA stability and folding— it is interesting that Sla1 can inhibit access of pre-tRNAs to Trm1. Since both factors promote tRNA function but Sla1 impedes Trm1, it would thus be interesting to speculate about possible situations (stresses, cellular growth conditions) in which a modification at G26 is of greater benefit relative to a tRNA that accesses La-associated RNA chaperone activity. It remains to be found whether a relationship between La and Trm1 exists in humans, where La proteins harbor an additional RNA recognition motif (RRM2 or xRRM) within a more divergent C-terminal domain that has been implicated in La’s RNA chaperone activity (50–52). The extent to which the C-terminal domain influences tRNA binding *in vivo* remains unknown, although its propensity for sequence-independent binding to structured, hairpin-containing RNA (51, 53) may affect how La engages the pre-tRNA body. Further, exploring how La may influence the installation of other modifications occurring at the pre-tRNA stage will likely be an active area of future research.

Another unexpected but related result concerns the link between apparent mature suppressor tRNA levels and tRNA-mediated suppression activity (16). We found that apparent increases in mature suppressor tRNA levels can at least be partially attributed to a decrease in modification at G26. As relative pre-tRNA levels are often quantitated using probes that anneal to the pre-tRNA intron, a probe that also overlaps with the nearby G26 base might then be reporting on a combination of both pre-tRNA abundance and G26 modification levels, supporting the idea of exercising caution when designing northern blotting probes to avoid sites of bulky modifications on the Watson-Crick face (29).

Finally, our work identifying nuclear and mitochondrial functions for Trm1 in *S. pombe* adds to the growing body of literature proposing coordination of the nuclear and mitochondrial genomes through tRNA modifications (54). In budding and fission yeast, this may be achieved by altering the balance of transcription of the nuclear– and mitochondrially-targeted Trm1 isoforms, which will in turn influence the degree of G26 dimethylation of nuclear– and mitochondrially-encoded tRNAs. Still, how Trm1-catalyzed tRNA modifications may alter cytoplasmic and mitochondrial translation remains unknown. Although we showed that there were no defects in bulk translation upon deletion of Trm1, we do not exclude the possibility that dimethylation at G26 alters the dynamics of codon recognition, as has been described for methylation of human mitochondrial serine and threonine tRNAs at position 32 by METTL8 (55). Additionally, studying the interplay between Trm1 and other nuclear-encoded enzymes that similarly modify mitochondrial tRNAs will continue to inform our understanding of the control and potential co-regulation of the nuclear and mitochondrial genomes.

Our work underscores the complexity of tRNA maturation in a model eukaryote by highlighting the multi-functional nature of a tRNA modification enzyme. The data presented here support the growing idea that tRNA modification enzymes may have roles beyond catalysis, and that this holds true in prokaryotes and eukaryotes. The extent to which this applies to other tRNA modification enzymes, and RNA modification enzymes more generally, will be an exciting area of future research that will continue to inform our understanding of the mechanisms underlying RNA fold and function.

### Experimental Procedures

#### Yeast strains and constructs

Standard laboratory techniques were used to culture *S. pombe* cells at 32°C. Tag integration and knockouts were generated with a previously described PCR-based strategy and verified by PCR and western blotting (56) (strains are provided in supplementary table 1). For plasmid-mediated expression of the Trm1 isoforms, the coding sequences of M1 or M24 Trm1 were cloned into the pRep82X yeast expression vector (57). Mutations were introduced by site-directed mutagenesis and verified by sequencing (supplementary table 2). Plasmids were introduced with lithium acetate and selected on minimal media lacking supplements (EMM-ura) (58). tRNA-mediated suppression growth assays were performed as described (34).

#### RNA and protein extractions and northern and western blotting

*S. pombe* cells were grown at 32°C and harvested at an OD_600 nm_ of 0.6. Total RNA was isolated with hot acid phenol and northern analysis was performed as described using 15% TBE-urea polyacrylamide gels (34). Band intensities were quantified with ImageQuant TL software. For PHA26 northerns (21), relative modification index values represent the intensity of the TUC signal divided by the intensity of the PHA26 signal and normalized to a wild type strain or empty vector. For tRNA-mediated suppression northerns, blots were probed with the indicated ^32^P-labeled probes and an equimolar amount of unlabeled competitor probe to prevent hybridization of the labeled probe with the endogenous tRNA Ser^UGA^, as per (16) (probe sequences are provided in supplementary table 3). Total protein was extracted according to (34) and western blots were probed with a custom anti-Trm1 antibody (gift from Dr. Richard Maraia, NIH) at 1:2000, β-actin (abcam, ab8226) at 1:1250, and human La/SSB (abcam, ab75927) at 1:1000.

#### cDNA synthesis and qRT-PCR

1 μg Turbo DNase-treated RNA was reverse transcribed with the iScript cDNA synthesis kit (BioRad, 1708890), treated with 0.5 μL Rnase cocktail (Invitrogen, AM2286), and diluted 1:10 before quantification using the SensiFAST SYBR No-Rox kit (Bioline, BIO-98005). qRT-PCR was performed with 1 μM of each primer (a common reverse primer for both Trm1 isoforms and isoform-specific forward primers, probes provided in supplementary table 3) with the following qRT-PCR settings: 95°C for 10 minutes and 40 cycles consisting of 10 seconds at 95°C, 20 seconds at 60°C, and 20 seconds at 72°C. Trm1 levels were normalized to *act1* mRNA and normalized M1 Trm1 signal was subtracted from normalized M24 Trm1 signal to calculate relative M24 Trm1 levels.

#### Pulse labeling of mitochondrial protein synthesis

Pulse labeling was performed according to (59). Briefly, *S. pombe* cells were initially grown in EMM-ura + 2% glucose, then inoculated into fresh EMM-ura with 0.1% glucose and 2% galactose and grown at 32°C for 6 generations to a final OD_600_ less than 1 x 10^7^ cells/mL. Cell pellets corresponding to ∼2.5 x 10^7^ cells were washed with 500 μL reaction buffer (40 mM potassium phosphate pH 6.0, 2% galactose, 0.1% glucose), pelleted, resuspended in 500 μL reaction buffer with 10 mg/mL cycloheximide, and incubated at room temperature for 15 minutes. Cycloheximide was omitted to evaluate cytoplasmic translation. 5 μL [^35^S]-methionine was directly added to the cell suspension, mixed thoroughly, and incubated for 30 minutes at room temperature. Cells were pelleted and the pellet was resuspended in 75 μL solubilization buffer (1.8 M NaOH, 1 M β-mercaptoethanol, 0.01 mM PMSF) and mixed. 500 μL water was added and proteins were TCA-precipitated, followed by separation on a 17.5% SDS gel, transfer to a nitrocellulose membrane, and exposed to a Phosphor screen overnight.

#### Recombinant protein purification and electrophoretic mobility shift assay (EMSA)

Recombinant His-tagged protein expression was induced in *Escherichia coli* with 1 mM IPTG at 16°C for 18 hours and purified over a Ni^2+^ column (His-TRAP, GE-Amersham), followed by a second round of purification over a heparin column (Hi-TRAP, GE-Amersham). Proteins were concentrated into 1X EMSA buffer (20 mM Tris HCl pH 7.6, 100 mM KCl, 0.2 mM EDTA pH 8.0, 1 mM DTT) and quantified by SDS-PAGE (Figure 5). EMSAs were performed as described (18). Briefly, 3000 cpm PAGE purified, T7-transcribed ^32^P α-ATP-labeled pre-tRNAs (sequences provided in supplementary table 4) were heated to 95°C and slow-cooled to room temperature before addition to a 20 uL reaction containing 1X EMSA buffer. Increasing amounts of recombinant Trm1 were added to the reaction mix, followed by incubation at 32°C for 20 minutes. Reactions were cooled on ice for 2 minutes and complexes were resolved on 8% non-denaturing polyacrylamide gels run at 4°C and 100V. The proportion of bound tRNA (the sum of the initial binding event and any supershifts) was quantified with ImageQuant TL software and binding curves were fit to a non-linear specific binding curve (Specific binding with Hill slope with least square fit, using the equation Y=B_max_*X^h^/(K_D_^h^+X^h^, where B_max_ is the maximum specific binding and h is the Hill slope) with GraphPad Prism 10.0.

#### *In vitro* methylation and primer extension

5 μM T7-transcribed pre-tRNA Ser^UGA^ was methylated for 3 hours at 32°C in a 25 μL reaction containing 100 mM Tris HCl pH 7.5, 0.1 mM EDTA pH 8.0, 10 mM MgCl_2_, 40 mM NH_4_Cl_2_, 1 mM DTT, 1.28 mM SAM (NEB, B9003S), and 5 μM or 5μg recombinant Trm1. For competition between La and Trm1, 5 μM pre-tRNA Ser^UGA^ was pre-incubated alone or with 5 μM recombinant hLa or Sla1 in a 50 μL reaction for 20 minutes at 32°C in methylation reaction buffer without MgCl_2_ and SAM. Reactions were snap chilled on ice to preserve complexes, followed by the addition of 5 μM recombinant Trm1, 2 mM MgCl_2_, and 1.28 mM SAM. Reactions were incubated at 32°C and samples were removed at indicated time points and immediately purified by phenol: chloroform: isoamyl alcohol (25:24:1) extraction. Primer extensions were performed with SuperScript III (Invitrogen, 18080093) following standard methods. For time course reactions, quantifications of G26 modification at each time point were fit to a non-linear specific binding curve (One-phase association with least squares fit) in GraphPad Prism 10.0 with the equation Y=Y_0_ + (Plateau-Y_0_)*(1-exp(-K*X).

#### FRET assays

FRET assays were performed as described (17, 41). Briefly, unlabeled and Cy5– and Cy3-labeled RNA substrates were synthesized by IDT (sequences provided in supplementary table 4) and used at a final concentration of 25 nM. Where applicable, recombinant proteins were added to 400 μL reactions at a final concentration of 100 nM immediately prior to taking measurements. Fluorescence emission at 590 and 680 nm was recorded on a Cary Eclipse fluorimeter with readings taken in half-second time-points over a period of 180 seconds. Strand dissociation measurements were initiated immediately following strand annealing with the addition of 1 μM unlabeled competitor RNA. Strand annealing and dissociation rate constants were determined by calculating a FRET index (emission at 680 nm divided by emission at 590 nm) over time and normalizing values between 0 and 1. Curves were fitted to an equation for one-phase association (strand annealing) or one-phase decay (strand dissociation) in Graphpad Prism 10.0.

## Data Availability

All data pertaining to this manuscript is included in the main manuscript and supporting information.

This article contains supporting information.

## Author Contributions

Jennifer Porat: Conceptualization, Investigation, Formal analysis, Methodology, Validation, Writing—original draft, Writing—review & editing. Ana Vakiloroayaei: Investigation, Formal analysis, Methodology, Writing—review & editing. Brittney M. Remnant: Validation, Writing— review & editing. Mohammadaref Talebi: Validation, Writing—review & editing. Taylor Cargill: Validation, Writing—review & editing. Mark A. Bayfield: Conceptualization, Writing—review & editing, Supervision, Funding acquisition.

## Acknowledgements

We thank Dr. Richard Maraia for the Trm1 antibody and *maf1Δ* yeast strain.

## References

1. Phizicky, E. M., and Hopper, A. K. (2010) tRNA biology charges to the front. Genes and Development. 24, 1832–1860

2. Porat, J., Kothe, U., and Bayfield, M. A. (2021) Revisiting tRNA chaperones: New players in an ancient game. RNA. 27, 543–559

3. Hopper, A. K. (2013) Transfer RNA post-transcriptional processing, turnover, and subcellular dynamics in the yeast Saccharomyces cerevisiae. Genetics. 10.1534/genetics.112.147470

4. Kadaba, S., Wang, X., and Anderson, J. T. (2006) Nuclear RNA surveillance in Saccharomyces cerevisiae: Trf4p-dependent polyadenylation of nascent hypomethylated tRNA and an aberrant form of 5S rRNA. RNA. 10.1261/rna.2305406

5. Kadaba, S., Krueger, A., Trice, T., Krecic, A. M., Hinnebusch, A. G., and Anderson, J. (2004) Nuclear surveillance and degradation of hypomodified initiator tRNA Met in S. cerevisiae. Genes and Development. 18, 1227–1240

6. Alexandrov, A., Chernyakov, I., Gu, W., Hiley, S. L., Hughes, T. R., Grayhack, E. J., and Phizicky, E. M. (2006) Rapid tRNA decay can result from lack of nonessential modifications. Molecular Cell. 10.1016/j.molcel.2005.10.036

7. M. Tasak, M. and E. M. Phizicky Initiator tRNA lacking 1-methyladenosine is targeted by the rapid tRNA decay pathway in evolutionarily distant yeast species. PLoS Genetics. 18, e1010215

8. T. De Zoysa and E. M. Phizicky Hypomodified tRNA in evolutionarily distant yeasts can trigger rapid tRNA decay to activate the general amino acid control response, but with different consequences. PLoS Genetics. 16, e1008893

9. Stefano, J. E. (1984) Purified lupus antigen la recognizes an oligouridylate stretch common to the 3′ termini of RNA polymerase III transcripts. Cell. 36, 145–154

10. Rinke, J., and Steitz, J. A. (1982) Precursor molecules of both human 5S ribosomal RNA and transfer RNAs are bound by a cellular protein reactive with anti-La Lupus antibodies. Cell. 10.1016/0092-8674(82)90099-X

11. Teplova, M., Yuan, Y. R., Phan, A. T., Malinina, L., Ilin, S., Teplov, A., and Patel, D. J. (2006) Structural basis for recognition and sequestration of UUUOH 3′ temini of nascent RNA polymerase III transcripts by La, a rheumatic disease autoantigen. Molecular Cell. 21, 75–85

12. Kotik-Kogan, O., Valentine, E. R., Sanfelice, D., Conte, M. R., and Curry, S. (2008) Structural Analysis Reveals Conformational Plasticity in the Recognition of RNA 3′ Ends by the Human La Protein. Structure. 16, 852–862

13. Bayfield, M. A., Vinayak, J., Kerkhofs, K., and Mansouri-Noori, F. (2019) La proteins couple use of sequence-specific and non-specific binding modes to engage RNA substrates. RNA Biology. 18, 168–177

14. Yoo, C. J., and Wolin, S. L. (1997) The yeast La protein is required for the 3’ endonucleolytic cleavage that matures tRNA precursors. Cell. 10.1016/S0092-8674(00)80220-2

15. Bayfield, M. A., and Maraia, R. J. (2009) Precursor-product discrimination by la protein during tRNA metabolism. Nature Structural and Molecular Biology. 16, 430–437

16. Huang, Y., Bayfield, M. A., Intine, R. V., and Maraia, R. J. (2006) Separate RNA-binding surfaces on the multifunctional la protein mediate distinguishable activities in tRNA maturation. Nature Structural and Molecular Biology. 13, 611–618

17. Naeeni, A. R., Conte, M. R., and Bayfield, M. A. (2012) RNA chaperone activity of human La protein is mediated by variant RNA recognition motif. The Journal of biological chemistry. 287, 5472–5482

18. Vakiloroayaei, A., Shah, N. S., Oeffinger, M., and Bayfield, M. A. (2017) The RNA chaperone La promotes pre-tRNA maturation via indiscriminate binding of both native and misfolded targets. Nucleic acids research. 45, 11341–11355

19. Copela, L. A., Chakshusmathi, G., Sherrer, R. L., and Wolin, S. L. (2006) The La protein functions redundantly with tRNA modification enzymes to ensure tRNA structural stability. RNA (New York, N.Y.). 12, 644–654

20. Ellis, S. R., Morales, M. J., Li, J. M., Hopper, A. K., and Martin, N. C. (1986) Isolation and characterization of the TRM1 locus, a gene essential for the N2,N2-dimethylguanosine modification of both mitochondrial and cytoplasmic tRNA in Saccharomyces cerevisiae. Journal of Biological Chemistry. 261, 9703–9709

21. Arimbasseri, A. G., Blewett, N. H., Iben, J. R., Lamichhane, T. N., Cherkasova, V., Hafner, M., and Maraia, R. J. (2015) RNA Polymerase III Output Is Functionally Linked to tRNA Dimethyl-G26 Modification. PLoS Genetics. 11, e1005671

22. Dewe, J. M., Fuller, B. L., Lentini, J. M., Kellner, S. M., and Fu, D. (2017) TRMT1-catalyzed tRNA modifications are required for redox homeostasis to ensure proper cellular proliferation and oxidative stress survival. Molecular and Cellular Biology. 37, MCB.00214-17

23. Ellis, S. R., Hopper, A. K., and Martin, N. C. (1987) Amino-terminal extension generated from an upstream AUG codon is not required for mitochondrial import of yeast N2,N2-dimethylguanosine-specific tRNA methyltransferase. Proceedings of the National Academy of Sciences of the United States of America. 84, 5172–5176

24. Suzuki, T. (2021) The expanding world of tRNA modifications and their disease relevance. Nature Reviews Molecular Cell Biology. 22, 375–392

25. Keffer-Wilkes, L. C., Soon, E. F., and Kothe, U. (2020) The methyltransferase TrmA facilitates tRNA folding through interaction with its RNA-binding domain. Nucleic Acids Research. 48, 7981–7990

26. Keffer-Wilkes, L. C., Veerareddygari, G. R., and Kothe, U. (2016) RNA modification enzyme TruB is a tRNA chaperone. Proceedings of the National Academy of Sciences of the United States of America. 113, 14306–14311

27. Johansson, M. J. O., and Byström, A. S. (2002) Dual function of the tRNA(m5U54)methyltransferase in tRNA maturation. RNA. 8, 324–335

28. Thodberg, M., Thieffry, A., Bornholdt, J., Boyd, M., Holmberg, C., Azad, A., Workman, C. T., Chen, Y., Ekwall, K., Nielsen, O., and Sandelin, A. (2019) Comprehensive profiling of the fission yeast transcription start site activity during stress and media response. Nucleic acids research. 47, 1671–1691

29. Khalique, A., Mattijssen, S., and Maraia, R. J. (2022) A versatile tRNA modification-sensitive northern blot method with enhanced performance. RNA. 28, 418–432

30. Jumper, J., Evans, R., Pritzel, A., Green, T., Figurnov, M., Ronneberger, O., Tunyasuvunakool, K., Bates, R., Žídek, A., Potapenko, A., Bridgland, A., Meyer, C., Kohl, S. A. A., Ballard, A. J., Cowie, A., Romera-Paredes, B., Nikolov, S., Jain, R., Adler, J., Back, T., Petersen, S., Reiman, D., Clancy, E., Zielinski, M., Steinegger, M., Pacholska, M., Berghammer, T., Bodenstein, S., Silver, D., Vinyals, O., Senior, A. W., Kavukcuoglu, K., Kohli, P., and Hassabis, D. (2021) Highly accurate protein structure prediction with AlphaFold. Nature. 596, 583–589

31. Ihsanawati Nishimoto, M., Higashijima, K., Shirouzu, M., Grosjean, H., Bessho, Y., and Yokoyama, S. (2008) Crystal Structure of tRNA N-2, N-2-Guanosine Dimethyltransferase Trm1 from Pyrococcus horikoshii. Journal of Molecular Biology. 383, 871–884

32. Motorin, Y., Muller, S., Behm-Ansmant, I., and Branlant, C. Identification of modified residues in RNAs by reverse transcription-based methods. Methods in Enzymology. 425, 21–53

33. Rijal, K., Maraia, R. J., and Arimbasseri, A. G. (2015) A methods review on use of nonsense suppression to study 3’ end formation and other aspects of tRNA biogenesis. Gene. 556, 35–50

34. Porat, J., and Bayfield, M. A. (2020) Use of tRNA-Mediated Suppression to Assess RNA Chaperone Function. in RNA Chaperones: Methods and Protocols (Heise, T. ed), pp. 107– 120, Springer US, New York, NY, 10.1007/978-1-0716-0231-7_6

35. Huang, Y., Intine, R. V., Mozlin, A., Hasson, S., and Maraia, R. J. (2005) Mutations in the RNA Polymerase III Subunit Rpc11p That Decrease RNA 3′ Cleavage Activity Increase 3′-Terminal Oligo(U) Length and La-Dependent tRNA Processing. Molecular and Cellular Biology. 25, 621–636

36. Iben, J. R., Epstein, J. A., Bayfield, M. A., Bruinsma, M. W., Hasson, S., Bacikova, D., Ahmad, D., Rockwell, D., Kittler, E. L. W., Zapp, M. L., and Maraia, R. J. (2011) Comparative whole genome sequencing reveals phenotypic tRNA gene duplication in spontaneous Schizosaccharomyces pombe la mutants. Nucleic Acids Research. 39, 4728– 4742

37. Boguta, M., Czerska, K., and Zoładek, T. (1997) Mutation in a new gene MAF1 affects tRNA suppressor efficiency in Saccharomyces cerevisiae. Gene. 185, 291–296

38. Lentini, J. M., Alsaif, H. S., Faqeih, E., Alkuraya, F. S., and Fu, D. (2020) DALRD3 encodes a protein mutated in epileptic encephalopathy that targets arginine tRNAs for 3-methylcytosine modification. Nat Commun. 11, 2510

39. Beenstock, J., Ona, S. M., Porat, J., Orlicky, S., Wan, L. C. K., Ceccarelli, D. F., Maisonneuve, P., Szilard, R. K., Yin, Z., Setiaputra, D., Mao, D. Y. L., Khan, M., Raval, S., Schriemer, D. C., Bayfield, M. A., Durocher, D., and Sicheri, F. (2020) A substrate binding model for the KEOPS tRNA modifying complex. Nat Commun. 11, 6233

40. Intine, R. V. A., Sakulich, A. L., Koduru, S. B., Huang, Y., Pierstorff, E., Goodier, J. L., Phan, L., and Maraia, R. J. (2000) Control of transfer RNA maturation by phosphorylation of the human La antigen on serine 366. Molecular Cell. 10.1016/S1097-2765(00)00034-4

41. Hussain, R. H., Zawawi, M., and Bayfield, M. A. (2013) Conservation of RNA chaperone activity of the human La-related proteins 4, 6 and 7. Nucleic Acids Research. 10.1093/nar/gkt649

42. Awai, T., Ochi, A., Ihsanawati, Sengoku, T., Hirata, A., Bessho, Y., Yokoyama, S., and Hori, H. (2011) Substrate tRNA recognition mechanism of a multisite-specific tRNA methyltransferase, Aquifex aeolicus Trm1, based on the X-ray crystal structure. Journal of Biological Chemistry. 286, 35236–35246

43. Marguerat, S., Schmidt, A., Codlin, S., Chen, W., Aebersold, R., and Bähler, J. (2012) Quantitative analysis of fission yeast transcriptomes and proteomes in proliferating and quiescent cells. Cell. 151, 671–683

44. Carpy, A., Krug, K., Graf, S., Koch, A., Popic, S., Hauf, S., and Macek, B. (2014) Absolute proteome and phosphoproteome dynamics during the cell cycle of Schizosaccharomyces pombe (Fission Yeast). Mol Cell Proteomics. 13, 1925–1936

45. Gogakos, T., Brown, M., Garzia, A., Meyer, C., Hafner, M., and Tuschl, T. (2017) Characterizing Expression and Processing of Precursor and Mature Human tRNAs by Hydro-tRNAseq and PAR-CLIP. Cell Reports. 20, 1463–1475

46. Belisova, A., Semrad, K., Mayer, O., Kocian, G., Waigmann, E., Schroeder, R., and Steiner, G. (2005) RNA chaperone activity of protein components of human Ro RNPs. RNA. 11, 1084–1094

47. Mayer, O., Rajkowitsch, L., Lorenz, C., Konrat, R., and Schroeder, R. (2007) RNA chaperone activity and RNA-binding properties of the E. coli protein StpA. Nucleic Acids Research. 10.1093/nar/gkl1143

48. Rajkowitsch, L., Chen, D., Stampfl, S., Semrad, K., Waldsich, C., Mayer, O., Jantsch, M. F., Konrat, R., Bläsi, U., and Schroeder, R. (2007) RNA chaperones, RNA annealers and RNA helicases. RNA Biology. 4, 118–130

49. Dewe, J. M., Whipple, J. M., Chernyakov, I., Jaramillo, L. N., and Phizicky, E. M. (2012) The yeast rapid tRNA decay pathway competes with elongation factor 1A for substrate tRNAs and acts on tRNAs lacking one or more of several modifications. RNA. 18, 1886– 1896

50. Kuehnert, J., Sommer, G., Zierk, A. W., Fedarovich, A., Brock, A., Fedarovich, D., and Heise, T. (2015) Novel RNA chaperone domain of RNA-binding protein la is regulated by AKT phosphorylation. Nucleic Acids Research. 10.1093/nar/gku1309

51. Brown, K. A., Sharifi, S., Hussain, R., Donaldson, L., Bayfield, M. A., and Wilson, D. J. (2016) Distinct Dynamic Modes Enable the Engagement of Dissimilar Ligands in a Promiscuous Atypical RNA Recognition Motif. Biochemistry. 55, 7141–7150

52. Bayfield, M. A., Yang, R., and Maraia, R. J. (2010) Conserved and divergent features of the structure and function of La and La-related proteins (LARPs). Biochimica et Biophysica Acta – Gene Regulatory Mechanisms. 1799, 365–378

53. Martino, L., Pennell, S., Kelly, G., Bui, T. T. T., Kotik-Kogan, O., Smerdon, S. J., Drake, A. F., Curry, S., and Conte, M. R. (2012) Analysis of the interaction with the hepatitis C virus mRNA reveals an alternative mode of RNA recognition by the human la protein. Nucleic Acids Research. 40, 1381–1394

54. Bohnsack, M. T., and Sloan, K. E. (2018) The mitochondrial epitranscriptome: the roles of RNA modifications in mitochondrial translation and human disease. Cellular and Molecular Life Sciences. 75, 241–260

55. Schöller, E., Marks, J., Marchand, V., Bruckmann, A., Powell, C. A., Reichold, M., Mutti, C. D., Dettmer, K., Feederle, R., Hüttelmaier, S., Helm, M., Oefner, P., Minczuk, M., Motorin, Y., Hafner, M., and Meister, G. (2021) Balancing of mitochondrial translation through METTL8-mediated m3C modification of mitochondrial tRNAs. Molecular Cell. 81, 4810–4825.E12

56. Bähler, J., Wu, J. Q., Longtine, M. S., Shah, N. G., McKenzie, A., Steever, A. B., Wach, A., Philippsen, P., and Pringle, J. R. (1998) Heterologous modules for efficient and versatile PCR-based gene targeting in Schizosaccharomyces pombe. Yeast. 10.1002/(SICI)1097-0061(199807)14:10<943::AID-YEA292>3.0.CO;2-Y

57. Maundrell, K. (1993) Thiamine-repressible expression vectors pREP and pRIP for fission yeast. Gene. 123, 127–130

58. Kanter-Smoler, G., Dahlkvist, A., and Sunnerhagen, P. (1994) Improved method for rapid transformation of intact Schizosaccharomyces pombe cells. BioTechniques

59. Gouget, K., Verde, F., and Barrientos, A. (2008) In vivo labeling and analysis of mitochondrial translation products in budding and in fission yeasts. Methods in Molecular Biology. 457, 113–124

