## Supplemental Figures for "Crosstalk between the tRNA methyltransferase Trm1 and RNA chaperone La influences eukaryotic tRNA maturation"

This document includes

5 supporting figures

4 supporting tables

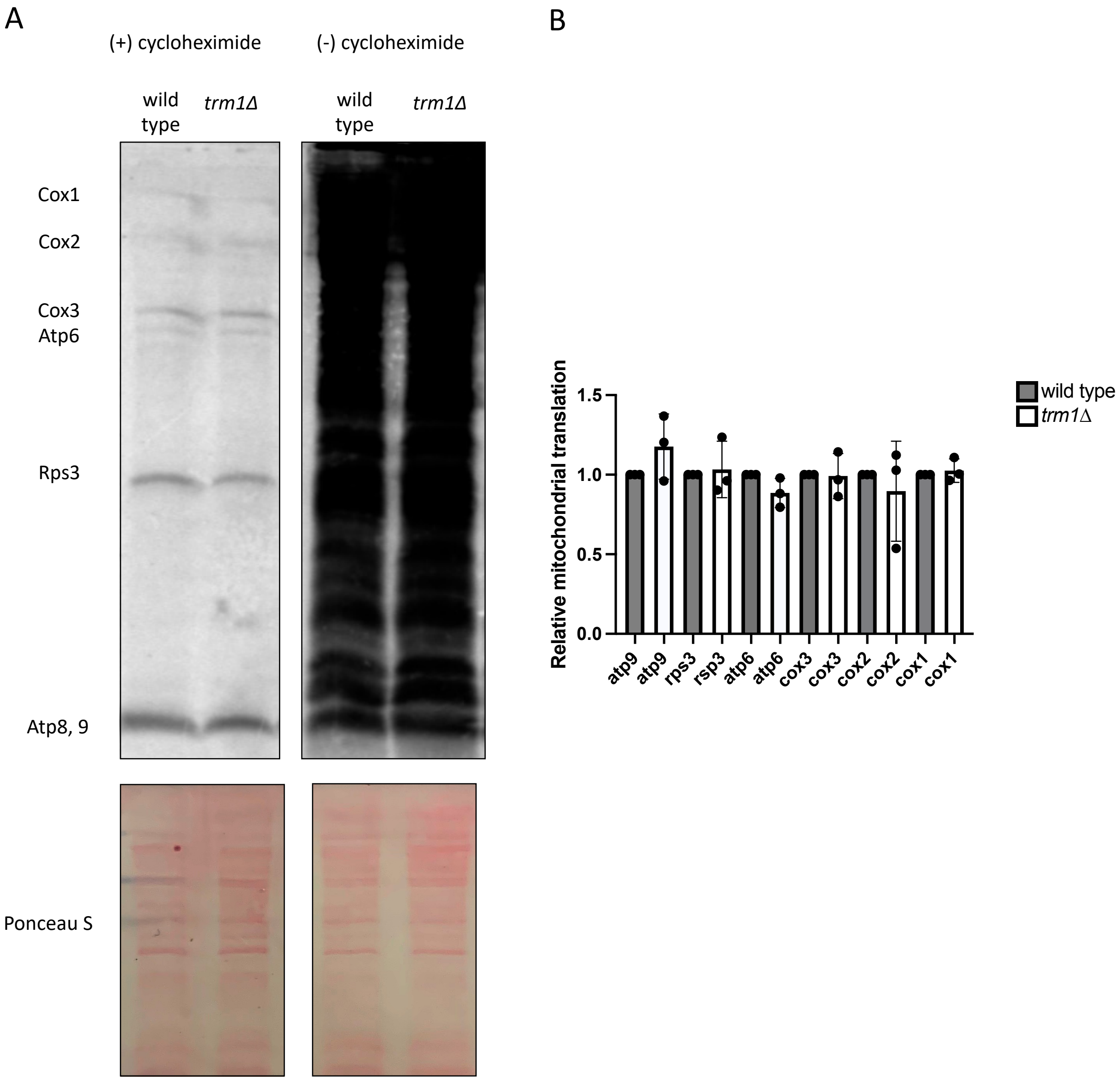


**Figure S1: Bulk cytoplasmic and mitochondrial translation are unaffected by Trm1 deletion**
30-minute pulse-labeling of newly synthesized mitochondrial (+ cycloheximide) and total (-cycloheximide) proteins in a wild type and *trm1Δ* strain. Mitochondrially synthesized proteins are indicated.

B) Quantification of mitochondrial translation products. Band intensity was normalized to total protein quantified by Ponceau S staining and normalized intensities were further normalized to a wild type strain (mean± standard error, n= 3 biological replicates).

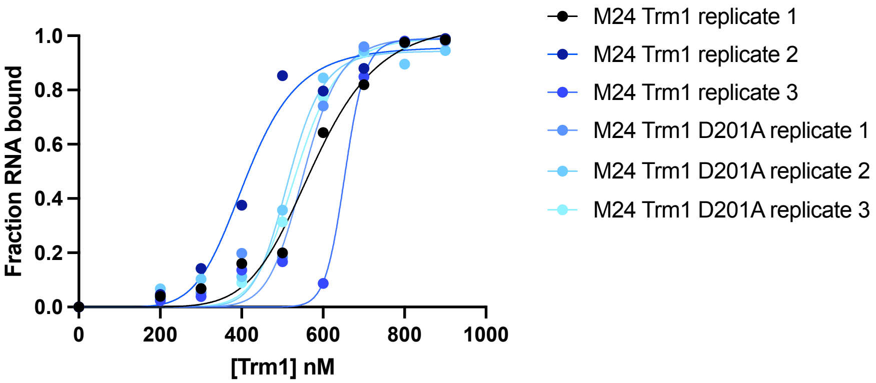


**Figure S2: Binding curves of M24 Trm1 and M24 Trm1 D201A EMSA technical replicates used to calculate $K_D$ values (associated with Figure 2).**

Curves were fit to a non-linear specific binding curve (specific binding with Hill Slope with least square fit, see methods for additional details).

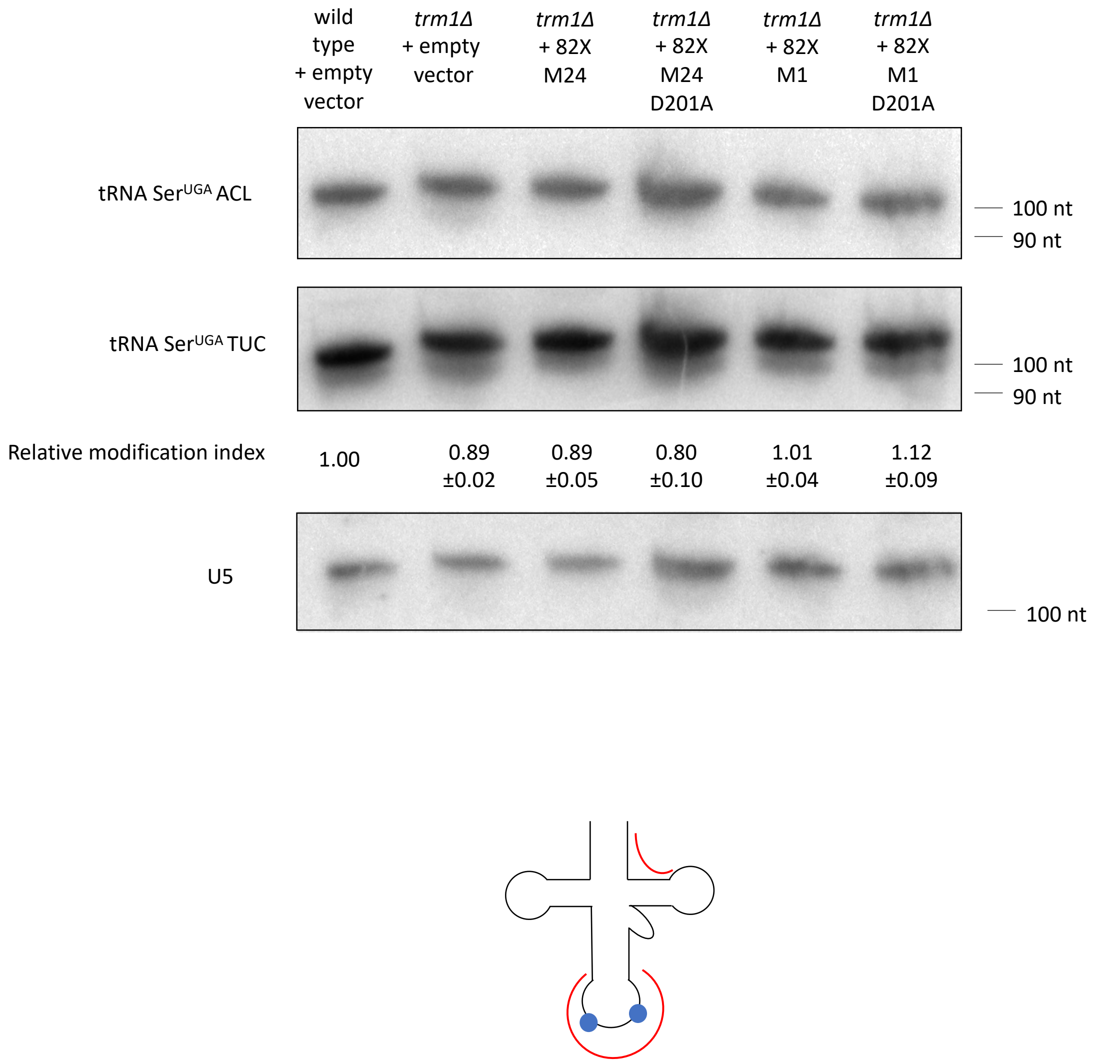


**Figure S3: Trm1 does not influence tRNA anticodon modifications that impair northern probe hybridization.**
PHA32/37 and TUC northern blots of endogenous tRNA Ser$^{UCA}$ in wild type and *trm1Δ* yeast strains overexpressing wild type or catalytically inactive Trm1 (mean± standard error, n= 3 biological replicates). Relative modification index was calculated as for with the PHA26 probe. Bottom: schematic depicting binding sites of the PHA32/37 and TUC probes, with modifications at position 32 (m$^3$C32) and 37 (t$^6$A37 and i$^6$A37) depicted with blue circles.

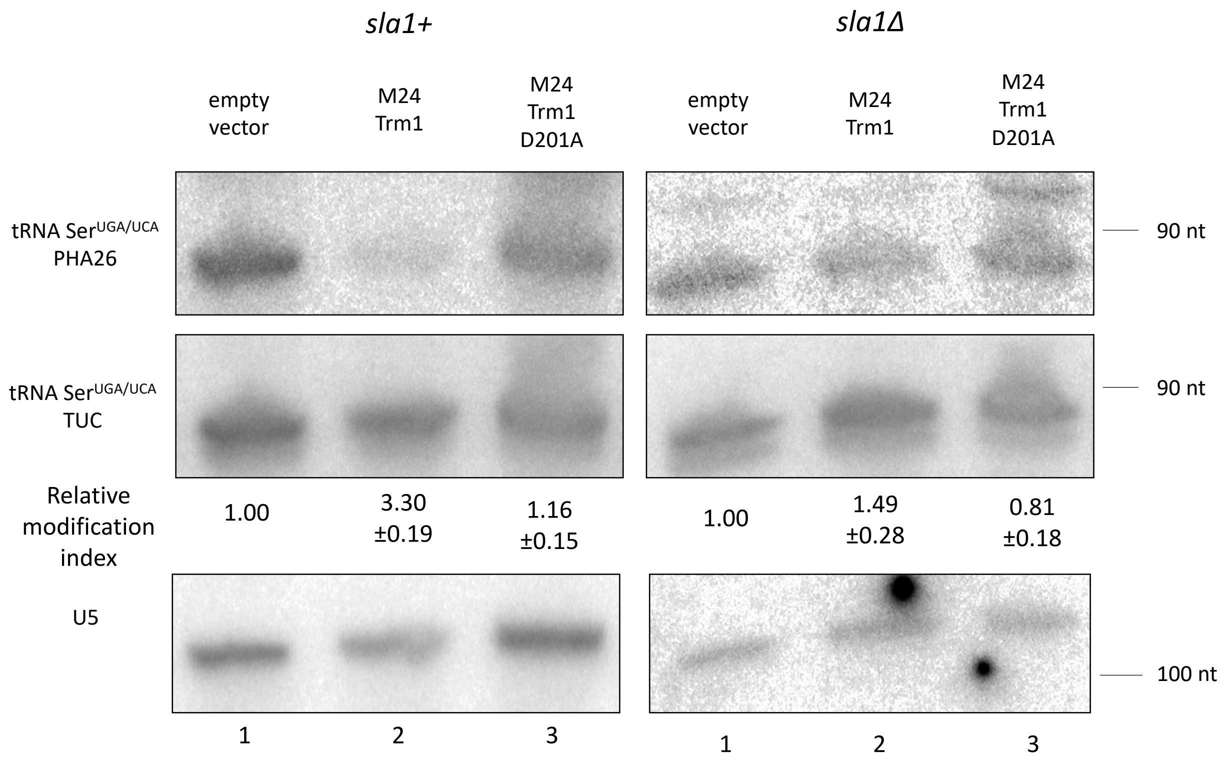


**Figure S4: tRNAs are Trm1-modified in the absence of Sla1.**
PHA26 and TUC northern blots of endogenous tRNA $Ser^{UCA}$ in a *sla1+* and *sla1Δ* strain overexpressing wild type or catalytically inactive Trm1 (mean± standard error, n= 3 biological replicates). Note that the PHA26 and TUC probes will also cross-react with the more lowly abundant suppressor tRNA MSer (*sla1+*) and $Ser^{UCA}$ (*sla1Δ*).

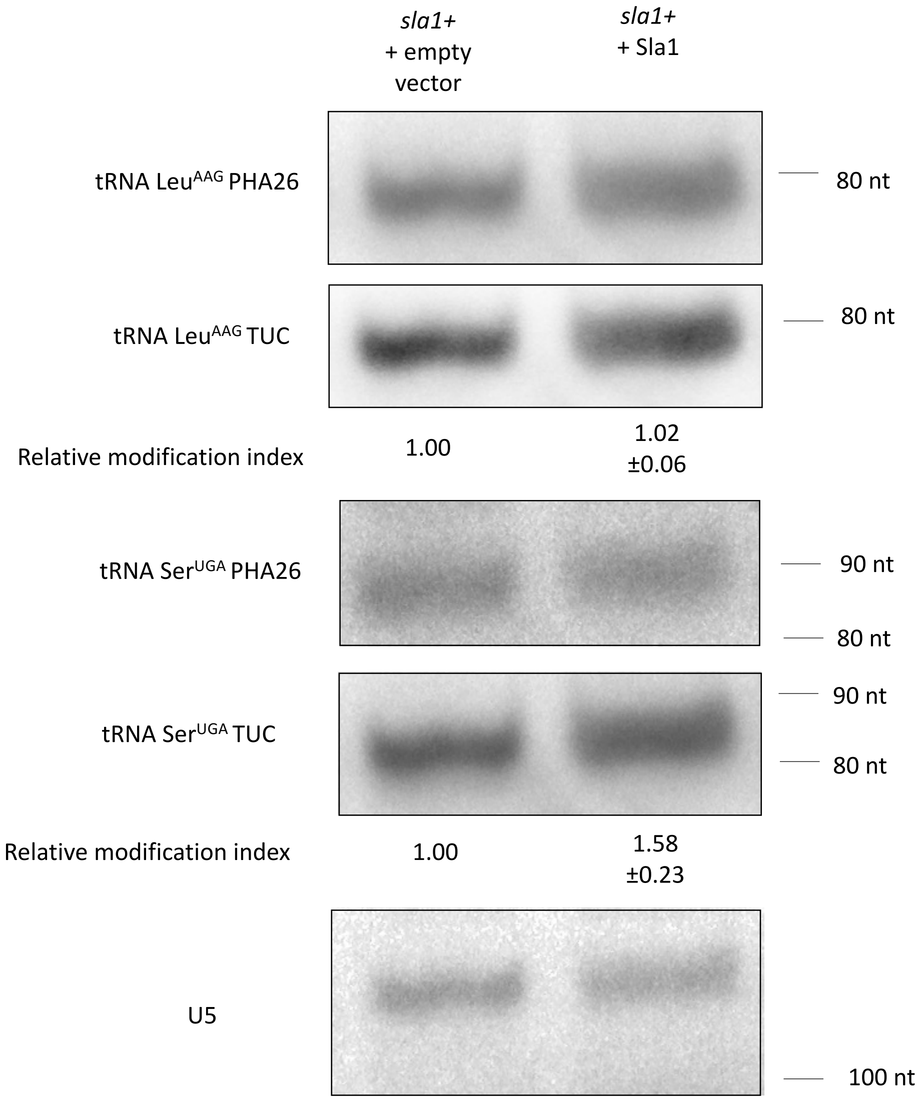


**Figure S5: Overexpressed Sla1 results in no further Trm1 modification defects in the context of endogenous Sla1**

PHA26 and TUC northern blots of tRNA Ser$^{UGA}$ and Leu$^{AAG}$ in a *sla1+* strain overexpressing Sla1 (mean± standard error, n= 3 biological replicates).

**Table S1: List of yeast strains used in this study**

| Figure | Strain | Description | Full genotype | Source |
|---|---|---|---|---|
| 1b, 1d, 5a, S1, S3, S4, S5 | y12088 | wild type | *h+ ura4-D18 leu1-32* | Lab stock |
| 1d, S1, S3 | yAV001 | *trm1Δ* | *h+ leu1-32 ura4 his7 lys1 trm1Δ::natMX6* | This study |
| 3a, 3b, 4a, 4c-e, 5a, 5b, 6, S5 | ySH9 | *sla1Δ* tRNA $Ser^{UCA}$ | *h- leu1-32::[tRNApSer7T-leu1+] sla1Δ::ura4-FOA+ ade6-704* | Huang *et al.,* 2005 |
| 3a, 3b, 4a, 4c-e, 5a | yJP039 | ySH9 *maf1Δ* | *h- leu1-32::[tRNApSer7T-leu1+] sla1Δ::ura4-FOA+ ade6-704 maf1Δ::kanMX6* | This study |
| 3a, 4b, 5a | yYH1 | tRNA MSer (C40U, U47.3C, C47.6U) | *h– leu1-32*::[*tRNAmSer7T-leu1+*] *ura4-D18 ade6-704* | Huang *et al.,* 2005 |
| 5a | yNB1 | *maf1Δ* tRNA Mser (C40U, U47.3C, C47.6U) | *h– leu1-32*::[*tRNAmSer7T-leu1+*] *ura4-D18 ade6-704 maf1Δ::kanMX6* | Arimbasseri *et al.,* 2015 |
| 3b | ySH18 | *maf1Δ* tRNA Mser (C40U, U47.3C, C47.6U) | *h- leu1-32::[tRNApSer7T U47:6leu1+] sla1Δ::ura4-FOA+ ade6-704* | Huang *et al.,* 2006 |

**Table S2: List of plasmids used in this study**

| Figure | Plasmid description |
|---|---|
| 1d, 3, 4, S3, S5 | pRep82X (empty vector) |
| 1d, 3, 4, S3, S5 | pRep82X + M24 Trm1 |
| 1d, 3, 4, S3, S5 | pRep82X + M24 Trm1 D201A |
| 1d, S3 | pRep82X + M1 Trm1 |
| 1d, S3 | pRep82X + M1 Trm1 D201A |
| 5b, 6, S4 | pRep4x (empty vector) |
| 5b, 6, S4 | pRep4X + Sla1 |
| 6 | pRep4X + hLa |
| 6 | pRep4X + hLa Q20A Y24A D33I |
| 6 | pRep4X + hLa Y114A F155A |
| 6 | pRep4X + hLa R142A R143A K148A K151A (Loop3) |
| 2, S2, 5b-d, 7 | pET28a + M24 Trm1 |
| 2, S2, 5d, 7 | pET28a + M24 Trm1 D201A |
| 5b-d, 7 | pET28a + hLa |
| 5b-d, 7 | pET28a + Sla1 |

**Table S3: List of northern blotting and qRT-PCR probes used in this study**

| Probe | Sequence |
|---|---|
| U5 | 5’ CTGGTAAAAGGCAAGAAACAGATACG 3’ |
| LeuN AAG PHA26 | 5’ AAACTAGCTCCATAACCACTCG 3’ |
| LeuN AAG TUC | 5’ AAGCTCGTGGGTTCGAGTCCCACCCCTTTCA 3’ |
| ThrM UGU PHA26 | 5’ GTACGCACTCTACCAATT 3’ |
| ThrM UGU TUC | 5’ GCTAATTAGAGGAATTGAACCTC 3’ |
| PheN GAA PHA26 | 5’ CAGTCTGTCATGCTC 3’ |
| PheN GAA TUC | 5’ TGTCACAAACCGGGATCGAACCGATGA 3’ |
| SerN UGA PHA26 | 5’ TTCAAGTCTAACTCCTTAACC 3’ |
| SerN UGA TUC (also used for primer extension) | 5’ GTCACCAGCAGGATTTGAACCTGCGCGG 3’ |
| SerN UGA ACL PHA | 5’ CGGCATTAGATTTCAAGTCTAA 3’ |
| ySH9 precursor | 5’ GAATACAGGATTGAAGTCT 3’ |
| ySH9 precursor competitor | 5’ GAATACAGGATTCAAGTCT 3’ |
| ySH9 mature | 5’ CCATTAGATTTGAAGTCTAA 3’ |
| ySH9 mature competitor | 5’ CCATTAGATTTCAAGTCTAA 3’ |
| ySH9 PHA26 | 5’ TGAAGTCTA ACTCCTT 3’ |
| ySH9 PHA26 competitor | 5’ TCAAGTCTAACTCCTT 3’ |
| LysN CUU precursor | 5’ CTTCTGATACCATTCGTAAGAGTC 3’ |
| Trm1 M1 qRT-PCR For | 5’ GTATGAAGCCAATATGCTAAGATTAAGTCAAAGG 3’ |
| Trm1 M24 qRT-PCR For | 5’ GCTTCTGCAAGTTTGACCGAG 3’ |
| Trm1 qRT-PCR Rev | 5’ CCGACCAAGCACGAATAGCTGTAACGCTTAAATCT 3’ |
| Act1 qRT-PCR For | 5’ TGCTCCTCCTGAGCGTAAATA 3’ |
| Act1 qRT-PCR Rev | 5’ CCGCTCTCATCATACTCTTGC 3’ |

**Table S4: List of RNA sequences used in this study**

| Name | Sequence |
|---|---|
| Pre-tRNA Ser UGA | 5’ GUCACUAUGUCCGAGUGGUUAAGGAGUUAGACUUGAAUCCUG UCUAGUCAUCUAAUGGGCUUUGCCCGCGCAGGUUCAAAUCCUGCU GGUGACGGUAUUU 3’ |
| 21F-Cy5 (top strand) | 5’ Cy5-AUGUGGAAAAUCUCUAGCAGU 3’ |
| 21R-Cy3 (bottom strand) | 5’ Cy3-ACUGCUAGAGAUUUUCCACAU 3’ |
| 21R (unlabeled bottom strand) | 5’ ACUGCUAGAGAUUUUCCACAU 3’ |
